# Ab initio side-chain sampling with PUD+ enables high-fidelity protein dynamics across AI-driven and classical simulations

**DOI:** 10.64898/2026.08.10.743906

**Authors:** Dianwei Wu, Tong Wang

## Abstract

The fidelity of molecular dynamics (MD) simulations fundamentally depends on the quality and coverage of the ab initio data used to parameterize the underlying force field, yet the role of side-chain conformational space remains insufficiently explored. In this study, we systematically investigate how comprehensive ab initio sampling of dipeptide conformations—specifically targeting side-chain degrees of freedom—impacts force field accuracy and MD simulation predictive power. We present the Protein Unit Dataset Plus (PUD+), a 40-million-conformation quantum mechanical dataset featuring unprecedented coverage of both backbone and side-chain conformational space. Machine learning force fields trained on PUD+ and integrated into AI^2^BMD simulations demonstrate superior energy and force prediction accuracy, capturing high-fidelity protein folding dynamics and the conformational flexibility of long-side-chain systems. Furthermore, leveraging PUD+ to reparameterize the CMAP term of the classical ff19SB force field markedly improves the description of intrinsically disordered protein (IDP) dynamics and IDP-ligand binding. Collectively, these results demonstrate that ab initio sampling of dipeptide side-chain conformations enables high-fidelity modeling of protein dynamics across both AI-driven and classical simulation paradigms.

## Main

In the post-AlphaFold era, the paradigm of structural biology is rapidly shifting from static snapshots toward dynamic conformational ensembles. Molecular dynamics (MD) simulation has been recognized as one of the most widely used computational approach for exploring the dynamic behavior of biomolecules^1,2^, offering critical insights into biological mechanism detection and molecular design. The fidelity of MD simulations, however, is fundamentally determined by the underlying force field: a well-parameterized force field calculates potential energies and atomic forces with high accuracy, which in turn enables simulation trajectories that faithfully capture the underlying conformational landscape.

To date, widely used classical MD simulations primarily rely on empirical force fields constructed using predefined analytical functional forms^3–5^. Despite the different functional details, such force fields typically require dipeptide datasets calculated at the density functional theory (DFT) level to derive well-defined parameters for bonded terms, which serve as a cornerstone that ultimately dictates the fidelity of the resulting simulations. In contrast, rapidly evolving AI-driven simulations employ machine learning force fields (MLFFs), which demand extensive DFT data for model training. Exemplified by our previous work AI^2^BMD^6^, we developed a fragmentation strategy to construct the Protein Universal Dataset (PUD), consisting of 20 million main-chain-sampling-enhanced conformations spanning the 20 canonical dipeptides for training the MLFF ViSNet^6^. It represents a major advance toward generalizable AI-driven MD simulations for biomolecular systems^7–9^.

While backbone dynamics are essential, protein side chains are the ultimate determinants of molecular recognition^10^, ligand binding^11,12^, and protein-protein interactions^13^, making their accurate modeling indispensable for elucidating functional mechanism in structural biology and designing molecules and drugs. Rotamer libraries derived either from statistical analysis of the Protein Data Bank (PDB), such as the Dunbrack backbone-dependent rotamer library^14^ or from dipeptide simulations, such as the MEDFORD rotamer library^15^, have been widely adopted into the parameterization workflows of classical force fields^3^. However, statistically derived libraries may introduce inherent conformational biases toward the specific secondary structures prevalent in the PDB^16^, whereas simulation-based approaches^15,17^ remain computationally prohibitive when attempting to comprehensively sample the full rotameric landscape of all 20 amino acids across diverse backbone contexts. This bottleneck is particularly pronounced in AI-driven simulations, where, on the one hand, deep neural networks with the millions to billions of parameters require vastly more training data. On the other hand, the combinatorial complexity of the side-chain conformational space renders exhaustive sampling and calculation at the ab initio level impractical for any target system. Consequently, the field has lacked a high-fidelity, quantum-accurate resource capable of refining both AI-driven and classical simulations at the full-atom, side-chain level.

In this study, we constructed the Protein Unit Dataset Plus (PUD+), a 40-million-conformation, DFT-level dipeptide dataset featuring comprehensive sampling of both side-chain and backbone conformations. Figure 1 illustrates the overall workflow of this study. We first assembled PUD+ by efficiently sampling and accurately calculating 20 million diverse side-chain conformations of the 20 canonical dipeptides and the ACE-NME, and integrating them with the original main-chain-sampling-enhanced PUD dataset introduced in AI^2^BMD. Conformational space analysis and structural clustering of both main-chain dihedrals φ and ψ and side-chain dihedrals χ1 and χ2 demonstrate that PUD+ explores a substantially broader structural diversity across both backbone and side-chain conformations compared to the original PUD and the traditional rotamer library-derived datasets. By training a series of MLFFs on varying subsets of PUD+, we achieved distinct performance gains driven by both the expanding scale of the dataset and the enhanced diversity of side-chain sampling. We then performed AI^2^BMD simulations with the MLFF trained on PUD+. Specifically, a complete protein folding process from a fully extended structure to a stable α-helix conformation was observed during the Ace-Ala_15_-Nme simulation. Furthermore, AI^2^BMD simulations of the STE7 docking motif using PUD+ yielded more accurate energy and force predictions, as well as more robust trajectories than those obtained using PUD. To examine the utility of PUD+ in classical MD simulations, we reparameterized the CMAP potential term of the Amber ff19SB force field and evaluated the resulting simulated dynamic behavior of both monomeric intrinsically disordered proteins (IDPs) and their ligand-bound complexes—a notorious challenge for conventional empirical force fields due to the high flexibility of IDPs and their sensitivity to backbone-sidechain coupling^18–21^. The ff19SB-PUD+ force field markedly improved the characterization of both IDP conformational dynamics and IDP-ligand binding, underscoring the value of the extensive conformational sampling provided by PUD+ in force field reparameterization. Collectively, our study demonstrates that the ab initio side-chain sampling encapsulated in PUD+ provides a comprehensive foundation for all-atom force field development, enabling high-fidelity, dynamic characterization of protein behaviors across both AI-driven and classical simulation paradigms.

**Figure 1.**
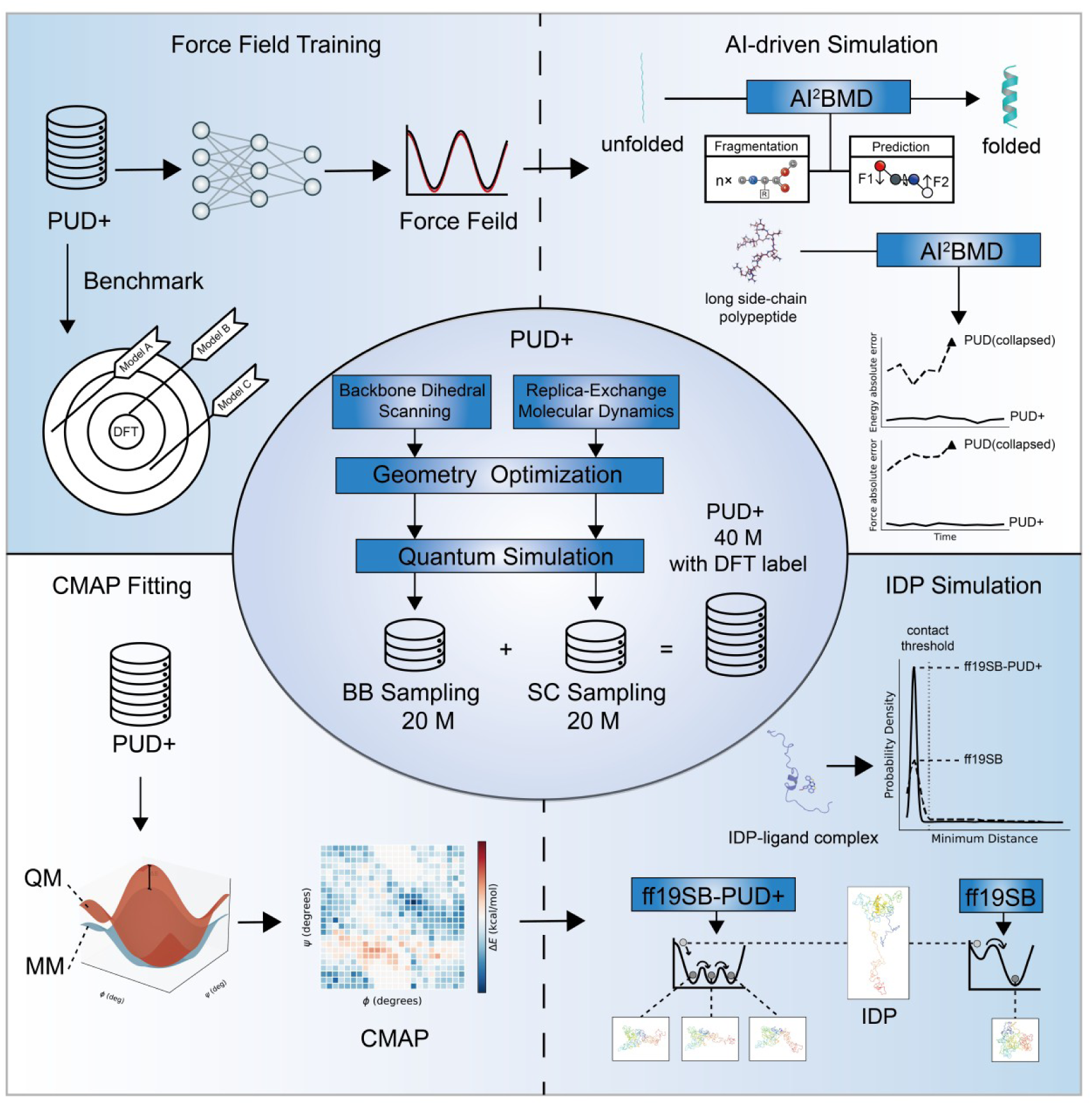
Schematic workflow of PUD+ dataset construction, force field design, and subsequent protein dynamics evaluation for both AI-driven and classical simulations. The 40-million-sample PUD+ dataset combines the original 20-million-sample backbone (BB) subset from AI^2^BMD, with a newly introduced 20-million-sample side-chain (SC) subset. For AI-driven simulations, we utilize PUD+ to train a series of MLFFs, which subsequently capture the complete protein folding events and robust dynamic behaviors in the corresponding AI^2^BMD simulations. For classical MD simulations, we reparameterize the CMAP term of the Amber ff19SB force field using PUD+ to simulate IDPs with high fidelity.

## Results

### Comprehensive sampling of dipeptide structural diversity

We constructed the complete PUD+ dataset by comprehensively sampling side-chain conformations (the SC sampling portion) and integrating them with the original PUD (the BB sampling portion). We conducted replica-exchange molecular dynamics (REMD)^22^ simulations and structural clustering of the 20 canonical protein dipeptides to obtain 5,000 representative structures spanning a wide distribution of side-chain conformations for each dipeptide. Then we performed quantum simulations at M06-2X/6-31g* level, a method adopted in AI^2^BMD^6^ to strike a favorable balance between accurate weak interaction profiling and acceptable computational cost^23^. Ultimately, a total of 20 million dipeptide conformations with comprehensively sampled side chains were obtained through the SC sampling workflow (see Methods for more details).

We then compared the main-chain Ramachandran plot (represented by φ and ψ dihedral angles) and the side-chain distribution plot (represented by χ1 and χ2 dihedral angles) between BB Sampling (the original PUD), SC sampling and the complete PUD+ dataset (Figure 2**a-f** for whole datasets, Supplementary Figure S1-S2 for each dipeptide type). Results demonstrated that BB sampling covered nearly the entire region around φ = 0° in the Ramachandran plot, which was the region of transient-state conformations (Figure 2**a**). SC sampling exhibited enhanced sampling of the steady-state conformations of various secondary structures (Figure 2**b**), aligning closely with the well-established empirical distributions^24^. The complete PUD+ dataset thus combined the advantages of both approaches, covering a wide region of transient-state conformations while accurately reproducing empirical distributions within the steady-state regions (Figure 2**c**). Regarding side-chain sampling distribution, BB sampling covered only 3 rotamers, (t, g-), (t, t), (t, g+) (Figure 2**d**), while SC sampling comprehensively captured all nine rotameric states within the side-chain χ1 and χ2 conformational space (Figure 2**e**). PUD+ inherited the superior side-chain coverage of SC sampling (Figure 2**f**). Consequently, we reasoned that integrating SC sampling with BB sampling was highly advantageous, combining their respective strengths to enable comprehensive exploration of both main-chain and side-chain conformational spaces.

**Figure 2.**
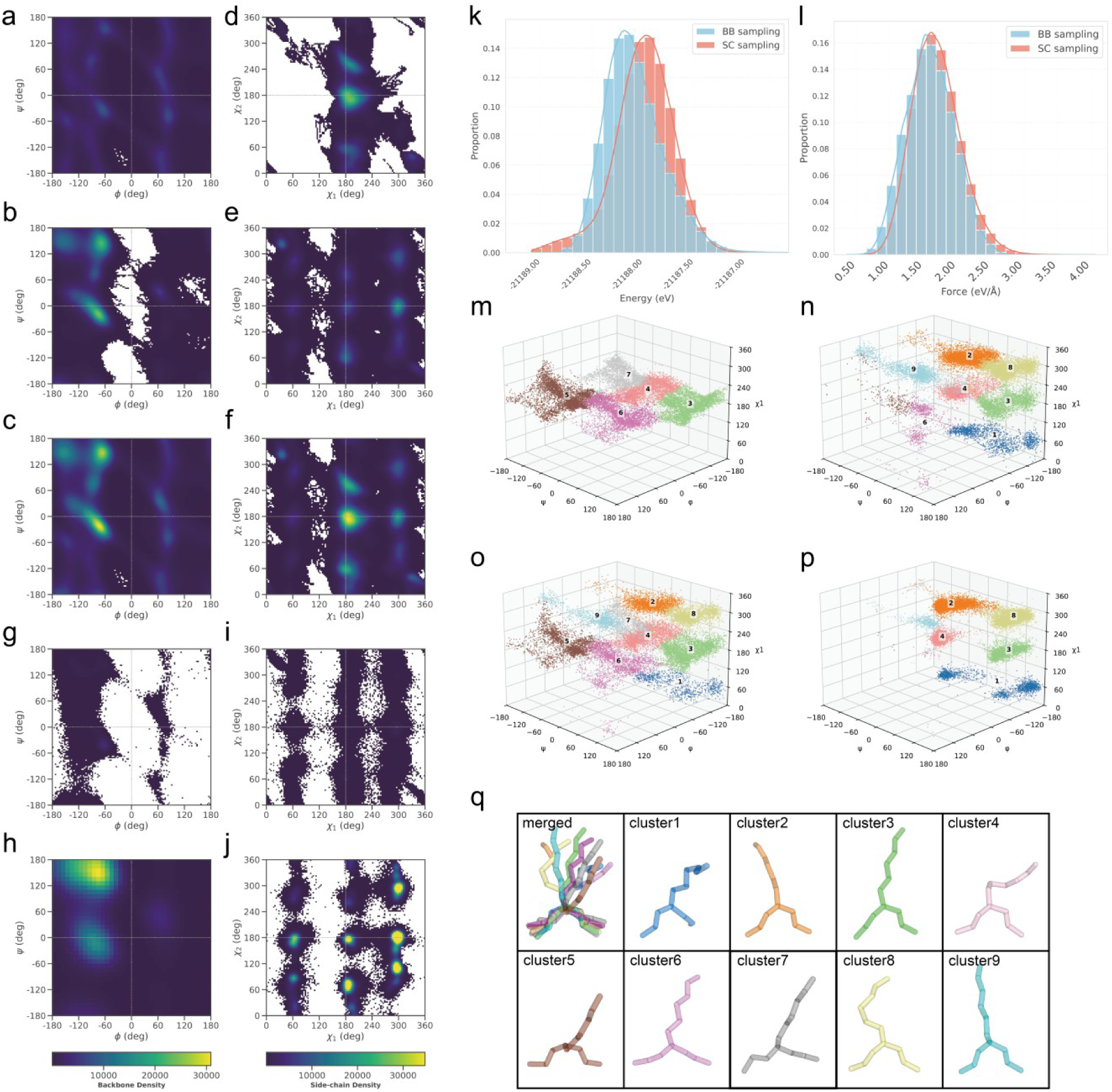
Conformational space coverage, energy-force distributions and structure clustering of PUD+ compared with other datasets. **a**-**c**, Ramachandran plots of main-chain dihedral angles φ, ψ derived from BB sampling (**a**), SC sampling (**b**), the complete PUD+ dataset (**c**). **d-f**, Distributions of side-chain dihedral angles χ1 and χ2 derived from BB sampling (**f**), SC sampling (**g**) and the complete PUD+ dataset (**h**). **g-j**, Ramachandran plot of main chain φ and ψ (**g**,**h**) and the distribution of side chain χ1 and χ2 (**i**,**j**) derived from the sidechain rotamer library Dunbrack 2010 (raw dataset) (**g**,**i**) and the MEDFORD library (**h**,**j**). **k**,**l**, Comparison of the distributions of potential energies (**k**) and atomic forces on side-chain atoms (**l**) derived from BB sampling and SC sampling, using the arginine dipeptide as a representative example. **m**-**p**, The structural clustering results of BB sampling (**m**), SC sampling (**n**), PUD+ (**o**) and Dunbrack raw dataset (**p**), using the arginine dipeptide as a representative example. The conformational space is defined by three dimensions: the backbone dihedral angles φ and ψ, and the first dihedral angle of side chain, χ1. The 9 clusters are visualized by different colors. **q**, The representative structures of each cluster.

We also compared PUD+ with rotamer library-based dipeptide datasets. The main-chain Ramachandran plot and side-chain χ1, χ2 distribution plot of the raw dataset used to generate Dunbrack backbone-dependent rotamer library (hereafter referred to as the Dunbrack raw dataset)^14^ and the Metadynamics of Dipeptides for Rotamer Distribution (MEDFORD) rotamer library^15^ are shown in Figure 2**g-j**. The main-chain Ramachandran plot of the Dunbrack raw dataset scarcely covered the transient-state region around φ = 0 (Figure 2**g**), while the MEDFORD rotamer library lacked key details in the main-chain Ramachandran plot due to the coarse binning of the main-chain dihedrals φ and ψ (Figure 2**h**), especially when compared to the continuous distributions in PUD+. Regarding the side-chain conformational space, similar to PUD+, both the Dunbrack raw dataset and the MEDFORD rotamer library captured all nine rotameric states, indicating the effectiveness of this approach for exploring the side-chain conformations (Figure 2**i-j**). Additionally, PUD+ uniquely outperformed these rotamer library-based datasets by providing full 3D coordinates, potential energies (Figure 2**k** for the arginine dipeptide) and atomic forces (Figure 2**l** for the arginine dipeptide) at DTF level, which is especially advantageous for training MLFFs. The potential energy and atomic force distribution plots for the remaining dipeptide types are shown in Supplementary Figure S3-S5.

To account for the coupling between the main-chain and side-chain dihedrals, we explored the 3D conformational space defined by main-chain dihedrals (φ and ψ) and the side-chain dihedral χ1 (Figure 2**m-p**), from which we identified nine representative structures, using the arginine dipeptide as a representative case (Figure 2**q**). As illustrated in Figure 2**m**, the BB sampling was confined to five discrete clusters and exhibited restricted exploration of the conformational space, scarcely sampling beyond the χ1 = 180° plane. In contrast, SC sampling achieved much more comprehensive coverage of the χ1 torsional space (Figure 2**n**). Furthermore, the merged PUD+ dataset distinctly enhanced sampling breadth, extending the conformational exploration seamlessly across the φ and ψ backbone dihedral angles and the χ1 side-chain axis (Figure 2**o**). Notably, the raw Dunbrack dataset exhibited sparse sampling in the φ > 0° region, yielding only five clusters (Figure 2**p**). The 3D conformational space analysis for each dipeptide type is shown in Supplementary Figure S6.

### Machine learning force fields trained on PUD+

To evaluate how side-chain enhanced-sampling improves the predictive accuracy of potential energies and atomic forces, we employed ViSNet-PIMA^25^, the improved version of ViSNet explicitly for long-range interaction modeling and trained the models on both the PUD+ and the original PUD datasets, respectively, using identical sample sizes (see Methods for details). We then assessed their performance on an independent PUD+ test set comprising samples completely unseen during model training (Figure 3**a**,**b**). The MLFF trained on PUD+ (termed the “PUD+ model”) markedly outperformed the one trained on PUD (termed the “PUD model”). The PUD+ model achieved a Mean Absolute Error (MAE) of 0.002576 kcal mol^-1^ per atom for potential energies and 0.05984 kcal mol^-1^ Å^-1^ for atomic forces. In comparison, the PUD model exhibited substantially higher errors, with an MAE of 0.01414 kcal mol^-1^ per atom for potential energies and 0.1825 kcal mol^-1^ Å^-1^ for atomic forces (Supplementary Table S1). Furthermore, we categorized the test set by side-chain properties to analyze the performance distribution in detail (Figure 3**c-j**, Supplementary Table S2 and S3). The PUD+ model maintained consistently low and stable MAEs across all dipeptide types. In contrast, the PUD model showed pronounced error spikes, particularly for dipeptides featuring long or charged side chains, indicating that the side-chain enhanced sampling mitigates prediction bias in these chemically challenging regions.

**Figure 3.**
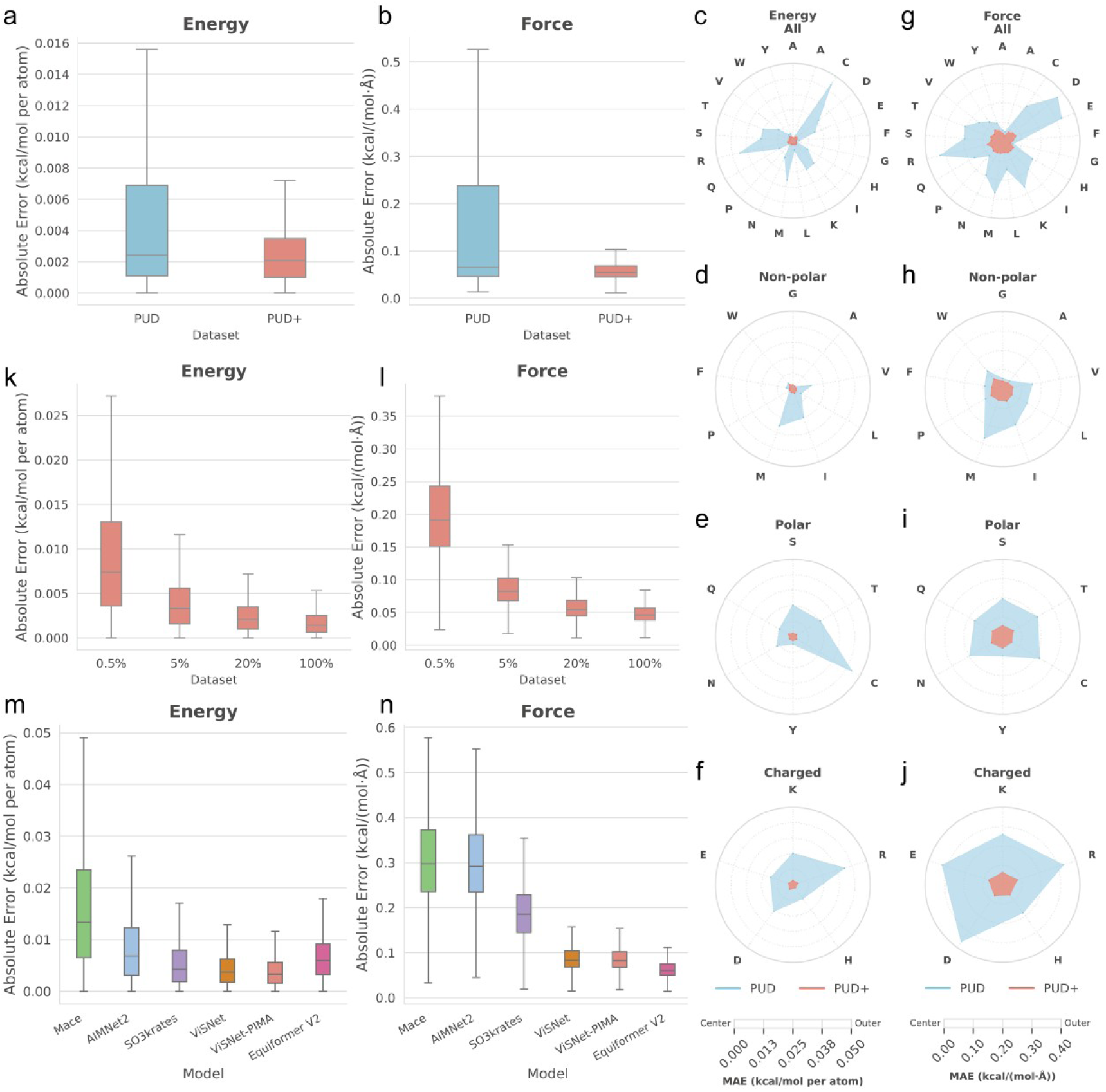
Evaluations of energy and force predictions by ViSNet-PIMA and other machine learning force fields trained on PUD+ dataset. 2 million samples were randomly selected from PUD+ dataset as the independent test set for all evaluations, while the remaining samples were employed for model training on PUD+. **a**,**b**, Absolute Error of potential energies (**a**) and atomic forces (**b**) predictions on the test set of PUD+. For a fair comparison, the ViSNet-PIMA models were trained on 7.6 million conformations either from the original PUD dataset (representing 40% of its total size) or PUD+ datasets (representing 20% of its total size), respectively. **c**-**j**, Radar plots show the MAE of potential energies (**c**-**f**) and atomic forces (**g**-**j**) across 20 dipeptides, corresponding to the results presented in **a** and **b**. These plots are categorized by side-chain properties: all 20 dipeptides (**c**, **g**), non-polar (**d**, **h**), polar (**e**, **i**), and charged groups (**f**, **j**). **k**,**l**, Absolute Error of potential energies (**k**) and atomic forces (**l**) on the test set, evaluated using the ViSNet-PIMA models trained on 0.5%, 5%, 20% and 100% samples of PUD+ training set, respectively. In **a**-**i**, skyblue represents ViSNet-PIMA trained on the PUD dataset, while salmon represents the model trained on the PUD+ dataset. **m**,**n**, Benchmark evaluations of potential energies (**m**) and atomic forces (**n**) on the PUD+ test set for different models trained on 5% samples of PUD+ training set (about 1.9 million samples).

To explore the dataset scaling behavior, we further trained ViSNet-PIMA on varying proportions of the PUD+ dataset, ranging from 0.5% to 100%, and evaluated each model on the independent test set (Figure 3**k**,**l**; see Methods for details). Our results reveal a clear scaling relationship between training set size and predictive accuracy: as the training volume increased, the MAEs for both potential energies and atomic forces decreased monotonically. Specifically, the model trained on 0.5% of the dataset yielded MAEs of 0.01075 kcal mol^-1^ per atom for potential energies (Figure 3**k**) and 0.2081 kcal mol^-1^ Å^-1^ for atomic forces (Figure 3**l**), which substantially decreased to 0.001850 kcal mol^-1^ per atom and 0.05024 kcal mol^-1^ Å^-1^, respectively, at full scale (100% of the dataset) (Supplementary Table S4). These findings confirm that the dataset scale is a primary determinant of MLFF prediction accuracy.

Given its large scale, extensive conformational coverage, balanced performance across all dipeptides, and high-accuracy DFT-level labels for potential energies and atomic forces, the PUD+ dataset serves as a robust benchmark for evaluating machine learning force fields (MLFFs) in biomolecular applications. To demonstrate this, we evaluated several state-of-the-art MLFFs using PUD+ as a benchmark (Figure 3**m** and 3**l**). All models were trained on 5% of the PUD+ training set and subsequently evaluated on the same independent test set for a fair comparison. For potential energy predictions, ViSNet-PIMA achieved the lowest average MAE of 0.004107 kcal mol^-1^ per atom, followed by ViSNet (0.004524 kcal mol^-1^ per atom) (Figure 3**m**, Supplementary Table S5). Regarding atomic force predictions, Equiformer V2 attained the lowest average MAE of 0.06550 kcal mol^-1^ Å^-1^, followed by ViSNet-PIMA (0.08977 kcal mol^-1^ Å^-1^) (Figure 3**n**). Collectively, these benchmarks highlight that high-quality, extensive conformational sampling is essential for developing highly accurate MLFFs for biomolecules.

### AI^2^BMD simulations of polypeptides with PUD+

To evaluate the practical efficacy of the PUD+ dataset for AI-driven MD simulations, we employed the ViSNet-PIMA model trained on PUD+ within the AI^2^BMD simulation framework to study the folding dynamics of the Ace-Ala_15_-Nme polypeptide. Through a 250 ps simulation in vacuum, the complete folding process, transitioning from an initial extended structure (0 ps) through a 3_10_-helix intermediate (120 ps) to a stable α-helix (220 ps) was faithfully captured, as evidenced by the RMSD trajectory evolution and representative snapshots in Figure 4**a**. This conformational evolution was further corroborated by the secondary structure fraction analysis (Figure 4**b**). We benchmarked the predictive accuracy of AI^2^BMD against a classical molecular mechanics (MM), Amber ff19SB. Using frames sampled at 10 ps intervals, we calculated reference potential energies and atomic forces at the DFT level. AI^2^BMD simulation demonstrated superior performance, achieving MAEs of 0.05118 kcal mol^-1^ per atom for relative potential energy (Figure 4**c**) and 1.7211 kcal mol^-1^ Å^-1^ for atomic forces (Figure 4**d**). In contrast, Amber ff19SB exhibited substantially higher errors, with 0.9473 kcal mol^-1^ per atom and 23.6177 kcal mol^-1^ Å^-1^ for relative energy and forces, respectively.

**Figure 4.**
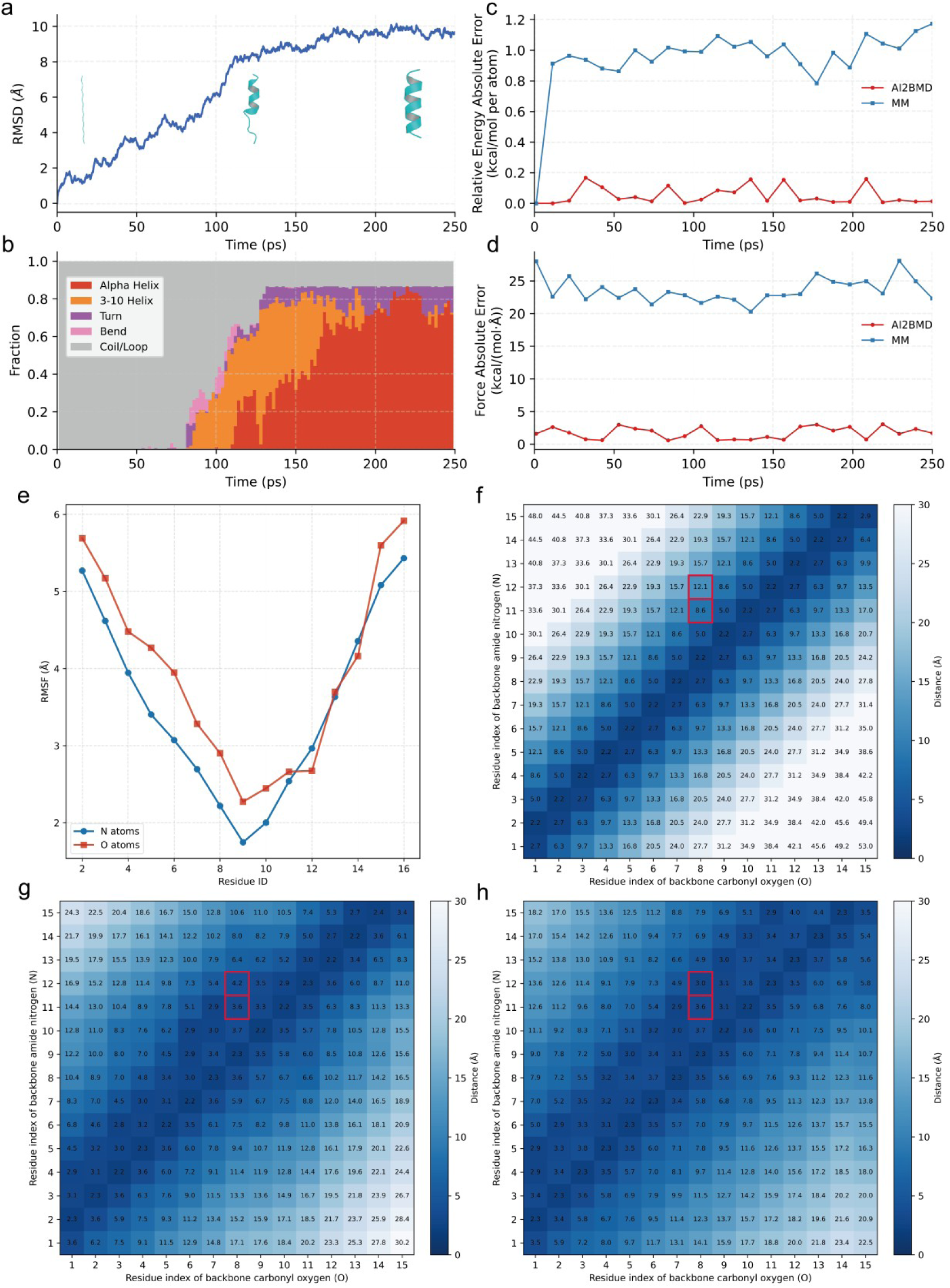
Analysis of Ace-Ala_15_-Nme folding process from AI^2^BMD simulation with PUD+ dataset. A 250 ps AI^2^BMD simulation in vacuum is conducted for Ace-Ala_15_-Nme, using ViSNet-PIMA model trained on the PUD+ dataset. The initial structure is a full extended structure. **a**, RMSD analysis of the 250 ps trajectory aligned to the initial frame, with representative structures shown at 0 ps (initial structure), 120 ps (intermediate structure), and 220 ps (final folded structure). **b**, Analysis of secondary structure evolution through the simulation trajectory. Five kinds of secondary structures that were formed during the simulation are shown with different colors. The other classes of secondary structures are omitted. **c**,**d**, The per atom relative absolute errors of the potential energies (**c**) and the atomic forces (**d**) during the whole simulation. Reference values were obtained via DFT calculations by ORCA 6.1.1 for the evenly sampled conformations from the trajectory. The relative energy is defined as the potential energy of each structure with that of the initial structure subtracted. For comparison, the energies and forces of evenly selected snapshots were also calculated by Amber ff19SB force field (denoted as “MM”). **e**, RMSF analysis of the Ace-Ala_15_-Nme simulation trajectory. The RMSFs of backbone carbonyl oxygen (O) and the amide nitrogen (N) of each residue are shown with red and blue curves, respectively. **f**-**h**, Residue-residue minimum distance maps for the conformation at 0 ps (initial state, **f**), 120 ps (intermediate state, **g**), and 220 ps (folded state, **h**). The red rectangles highlight the distances between backbone carbonyl oxygen (O) of 8th residue and the amide nitrogen (N) of 11th and 12th residues, respectively, differentiating the characteristic structural features of the 3₁₀-helix (**g**) and the α-helix (**h**).

We then analyzed the RMSF values of Ace-Ala_15_-Nme over the 250 ps simulation (Figure 4**e**), along with the distances between backbone amide nitrogens and backbone carbonyl oxygens across different conformational states (Figures 4**f-h**). The RMSF profile revealed that Ace-Ala_15_-Nme folded progressively from both termini toward the central region. As the folding progressed, the average distance between the backbone amide nitrogen of residue i and the backbone carbonyl oxygen of residue i-3 dropped sharply from 8.60 Å in the initial extended conformation (Figure 4**f**) to 3.64 Å at 120 ps (Figure 4**g**), characteristic of 3_10_-helix formation. By 220 ps, the distance between the backbone amide nitrogen of residue i and the backbone carbonyl oxygen of residue i-4 further decreased to 3.07 Å, falling below the corresponding i to i-3 distance of 3.36 Å. Such a reversal signals a transition from 3_10_-helix to an α-helical conformation (Figure 4**h**). Taken together, these results demonstrate that the AI^2^BMD framework, powered by PUD+-trained ViSNet-PIMA, recapitulates the structural folding transitions of Ace-Ala_15_-Nme with high fidelity.

We further evaluated how side-chain sampling enhances simulation fidelity and trajectory stability by conducting explicitly solvated simulations of a long-side-chain polypeptide within the AI^2^BMD framework, using ViSNet-PIMA trained on PUD+ versus PUD. The polypeptide (sequence RRNLKGLNLNLH, PDB ID: 2B9H) is a docking motif derived from serine/threonine protein kinase STE7^26^, characterized by multiple long-side-chain amino acid residues. To obtain diverse initial structures, we performed a REMD simulation starting from a folded conformation and clustered 10 representative conformations from the trajectory based on RMSD. These conformations (Figure 5**a**) were categorized as folded, intermediate, or unfolded structures according to their radius of gyration (Rg) (see Methods for details). We then conducted a cumulative 10 ns of explicit-solvent simulations (10 trajectories, 1 ns each) within the AI^2^BMD framework using ViSNet-PIMA trained on PUD+ and PUD, hereafter referred to as the “PUD+ simulations” and “PUD simulations”, respectively.

**Figure 5.**
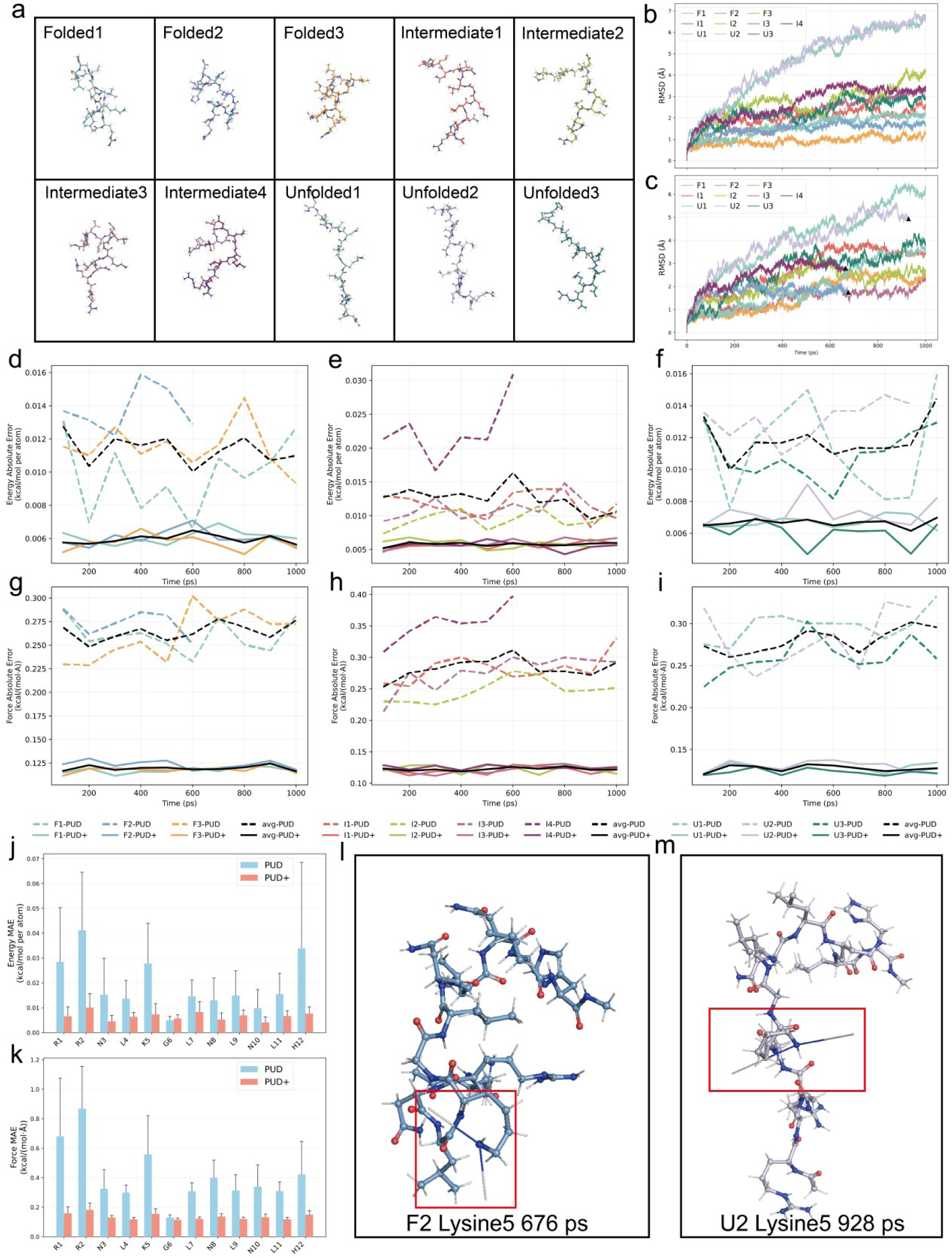
Analysis of protein dynamics for a long-side-chain polypeptide using AI^2^BMD simulation with PUD and PUD+ datasets in explicit solvent. The polypeptide is a docking motif derived from Serine/Threonine protein kinase STE7, characterized by the presence of multiple long-side-chain amino acid residues. A total of 20 ns of simulation was performed (20 trajectories of 1 ns each). The sampling covered 10 initial structural states (folded, intermediate, and unfolded) across two MLFFs trained on PUD and PUD+ datasets, respectively. **a**, 10 initial structures of the STE7 docking motif, including 3 folded structures, 4 intermediate structures and 3 unfolded structures. These structures were classified into the aforementioned categories according to their radius of gyration (Rg) (see Methods). **b**,**c**, RMSD analysis of the 10 trajectories (1 ns each) driven by the MLFFs trained on PUD+ (**b**) and PUD (**c**), respectively. All trajectories are aligned to the initial frame. In the legend, F, I, and U denote the folded, intermediate, and unfolded initial structures, respectively. The appended numbers indicate distinct representative structures within each category. The collapsed points are denoted by black triangles. **d-i**, The per atom absolute errors of the potential energies (**d**,**e**,**f**) and atomic forces (**g**,**h**,**i**). Panels from left to right display initial structures of folded (**d**,**g**), intermediate (**e**,**h**) and unfolded (**f**,**i**). Colors identify unique structures within each category, while dashed and solid lines represent trajectories driven by MLFFs trained on PUD and PUD+, respectively. Black lines represent the average MAE across all trajectories. Reference values were obtained via fragment DFT calculations by ORCA 6.1.1 for 10 evenly sampled conformations from each trajectory. For collapsed trajectories, only conformations sampled before collapsed point are included. **j**,**k**, Residue-wise, per atom MAEs of the potential energies (**j**) and the atomic forces (**k**). The same samples and reference values as in **d-i** were used to calculate the residue-wise MAE. The error bars in **j** and **k** indicate the standard deviations of 100 samples in PUD+ and 91 samples (with the collapsed frames excluded) in PUD. **l**,**m**, Ball-and-stick representations of two conformations at the collapse points, observed in the PUD trajectories. The collapsed atoms are highlighted with red rectangles.

The RMSD trajectories of the PUD+ and PUD simulations are shown in Figure 5**b**,**c**, respectively. All 10 PUD+ simulation trajectories successfully completed the full 1 ns simulation, whereas three PUD simulation trajectories—initiated from folded structure 2, intermediate structure 3, and unfolded structure 2-suffered simulation collapses at 676 ps, 665 ps, and 928 ps, respectively (indicated by black triangles in Figure 5**c**). To investigate the origin of this stability difference, we sampled frames from each trajectory at 10 ps intervals (excluding frames after the collapse points) and computed reference potential energies and atomic forces for dipeptide fragments at the DFT level (see Methods for details). Predictions were made by ViSNet-PIMA trained on PUD+ or PUD, and the resulting absolute errors in potential energies and atomic forces are presented in Figures 5**d-i**. Across all folded, intermediate, and unfolded trajectories, the PUD simulation exhibited substantially higher absolute errors, particularly in frames approaching the collapse points, whereas the PUD+ simulation maintained consistently low and stable errors, underscoring its enhanced simulation stability.

To pinpoint the source of these discrepancies, we further examined residue-wise, per-atom MAEs of potential energies (Figure 5**j**) and atomic forces (Figure 5**k**). The results revealed that the MAEs for long-side-chain residues (Arg1, Arg2, and Lys5) decreased most prominently in the PUD+ simulations, with other residues also showing error reductions to varying degrees. Figures 5**l**,**m** illustrate two PUD conformations at their respective collapse points, both of which exhibited severe unphysical distortions centered at the side-chain nitrogen atoms of Lys5, where the attached hydrogen atoms were ejected outward from their expected bonded positions. These findings collectively demonstrate that the extensive side-chain sampling afforded by PUD+ substantially advances the dynamic fidelity and the trajectory stability of proteins with long, flexible side-chains.

### Reparameterization CMAP potential in MM with PUD+

For molecular mechanics (MM) design and classical MD simulations, leveraging its high-quality data and extensive sampling of both backbone and side-chain conformations, PUD+ also provides an ideal foundation for CMAP reparameterization. We selected ff19SB^3^, a widely applied MM, that extends ff14SB by incorporating residue-specific CMAP potential energy corrections. To reparameterize the CMAP potential term of ff19SB using PUD+, we adopted a fitting procedure consistent with that originally employed for ff19SB^3^, ensuring full compatibility with the AMBER software suite. Furthermore, we implemented targeted optimizations to better align the resulting CMAP parameters with the conformational landscape and energy characteristics captured by PUD+ (see Methods for details).

Residue-specific CMAP parameters were fitted for each amino acid. We employed a 24 × 24 grid resolution, as used in Amber ff19SB^3^ force fields. For each grid point, the 10 nearest data points were aggregated, and their average potential energy was assigned to that point. The DFT-level potential energies in PUD+ were used to construct the DFT relative potential energy surface, whereas the same data points were evaluated using ff19SB without CMAP to obtain the corresponding MM relative potential energy surface. The final CMAP potential energy correction was obtained by subtracting the MM energy from the corresponding DFT energy at each (φ, ψ) grid point, and the full energy surface was then completed via bicubic spline interpolation between grid points, which is natively supported within the AMBER software suite. (Supplementary Figure S7 and S8).

### Simulations of IDPs using classical MD with a reparameterized CMAP

The accurate simulation of intrinsically disordered proteins (IDPs) remains a longstanding challenge for classical MD and the underlying molecular mechanics^21^. To evaluate ff19SB reparameterized with PUD+ (hereafter referred to as ‘ff19SB-PUD+’), we performed a 1,500 ns explicit-solvent simulation of the 526-residue IDP, Fused in Sarcoma (FUS) (Figure 6**a**), alongside a parallel 1,500 ns simulation using the original ff19SB as a control (see Methods for details).

**Figure 6.**
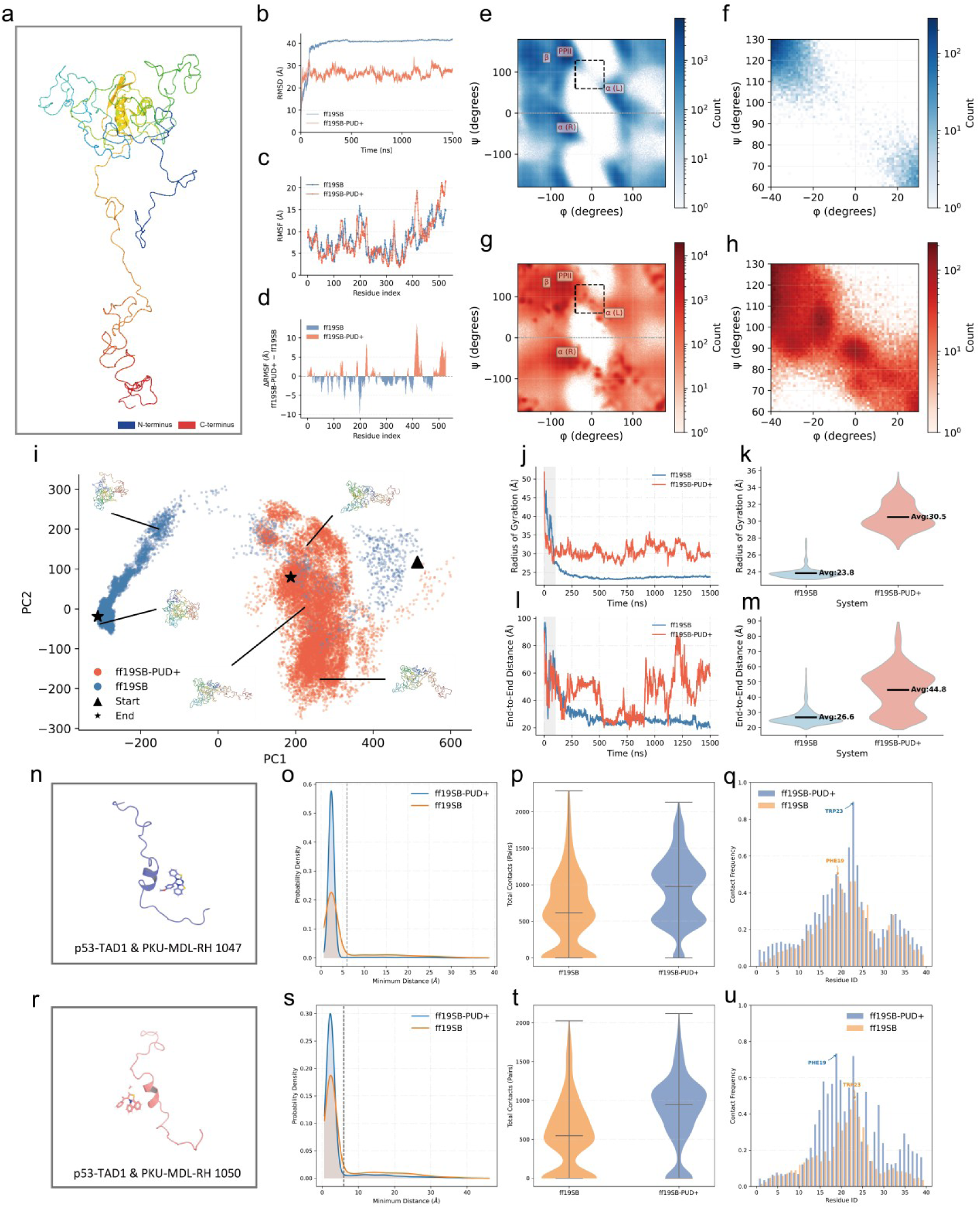
Analysis of intrinsically disordered protein (IDP) dynamics simulated with a PUD+ based CMAP potential. **a**-**e**, Simulations of the 526-reisdue IDP, Fused in sarcoma (FUS). **a**, The initial structure of FUS was obtained from Sarthak *el al*^21^. 1,500 ns simulations with explicit solvent were performed using the original Amber ff19SB force field and ff19SB with the PUD+ reparameterized CMAP energy (denoted as “ff19SB-PUD+”), respectively. **b-d**, RMSD (**b**), RMSF (**c**), and ΔRMSF (**d**) of the ff19SB and ff19SB-PUD+ trajectories; both trajectories were aligned to their initial frames. **e-h**, Ramachandran plots of the main-chain φ and ψ angles for the ff19SB trajectory (**e**, **f**) and the ff19SB-PUD+ trajectory (**g**, **h**). Panels **f** and **h** show zoomed-in views of the transition region connecting PPII and α(L), indicated by the black squares in e and g, respectively. The color bar indicates the number of points (counts) within each bin, where each point corresponds to the (φ, ψ) angle pair of a single residue in a single trajectory frame. **i**, PCA of the ff19SB and ff19SB-PUD+ trajectories based on the 3D coordinates of backbone Cα atoms. The starting frame is denoted by a black triangle and the ending frame by a black star. Conformations from both trajectories were clustered, yielding five clusters, with representative structures shown for each. **j**,**k**, Time series (**j**) and violin plots (**k**) of the radius of gyration (Rg). **l**,**m**, Time series (**l**) and violin plots (**m**) of the end-to-end distance (between N- and C-termini). The gray regions in **j** and **l** indicate the initial equilibration phase, which was excluded from the Rg distributions shown in the violin plots (**k**,**m**). **n**-**u**, Simulations of p53 transactivation domain 1 (TAD1) complexed with two compounds. Human p53 TAD1 (PDB ID, 2K8F) is a 39-residue IDP, which binds to the compounds PKU-MDL-RH 1047 (C_22_H_16_N_3_OS ^+^, denoted as “1047”) and PKU-MDL-RH 1050 (C_23_H_16_NOS ^+^, denoted as “1050”)^28^, respectively. MD simulations of p53-TAD1 complexed with the compound 1047 (**n**) and 1050 (**r**) were conducted by ff19SB and ff19SB-PUD+ for 3 μs each. **o**,**s**, Minimum distance distributions of the p53-TAD1 complexed with 1047 (**o**) and 1050 (**s**). Dashed lines indicate the contact threshold defined by a minimum distance less than 6 Å. **r**,**v**, Violin plots showing the number of contacts formed between p53-TAD1 and 1047 (**p**) or 1050 (**t**). Total contacts are defined as the total number of protein-ligand atom pairs within the distance cutoff of 6.0 Å. **q**,**u**, Per-residue contact frequencies of p53-TAD1-1047 (**q**) and p53-TAD1-1050 complexes (**u**). Contact frequency is defined as the fraction of total frames in which a given residue contacts with the compound.

We first assessed the overall structural stability and flexibility across the two trajectories using RMSD, RMSF, and ΔRMSF (Figures 6**b-d**). Compared to ff19SB-PUD+, the wild-type ff19SB control exhibited a larger overall RMSD, reflecting its transition from a relatively loose initial conformation to a compact folded state. However, the two systems differed markedly in their RMSD dynamics: the control rapidly reached and then plateaued at this elevated RMSD level with minimal further fluctuation, whereas ff19SB-PUD+ showed persistently large RMSD fluctuations throughout the trajectory, consistent with the protein remaining in a disordered state. ΔRMSF analysis revealed increased flexibility concentrated in the C-terminal tail region (residues 410-526), consistent with the conformational heterogeneity expected of a disordered chain. To further characterize backbone conformational preferences, we examined the Ramachandran distributions of φ and ψ angles for both trajectories (Figures 6**e-h**). While both force fields sampled the canonical polyproline II (PPII) and α(L) basins, ff19SB-PUD+ showed a markedly higher population density in the transition region connecting PPII and α(L) (Figures 6**f**, **h**), suggesting a more facile interconversion between these two backbone states. To assess the diversity of the global conformational ensemble, we performed principal component analysis (PCA) on the backbone Cα coordinates of both trajectories (Figure 6**i**). Clustering of the projected conformations identified five representative clusters for each trajectory. The ff19SB-PUD+ ensemble spanned a broader region of the principal component space, with representative clusters exhibiting more heterogeneous overall topologies compared to the more compact clusters observed for ff19SB, indicating that ff19SB-PUD+ samples a wider range of global conformations.

We next assessed global chain dimensions using the radius of gyration (Rg) and end-to-end distance, two metrics widely adopted in benchmark studies of IDP-targeted force fields^18,20,21^. As shown in Figures 6**j** and 6**k**, the ff19SB control group yielded substantially lower Rg values, reflecting its well-documented tendency to generate overly compact, globular conformations. In contrast, ff19SB-PUD+ produced consistently higher Rg values, indicating a more realistic representation of the extended and disordered ensemble characteristic of IDPs. This distinction was further supported by the end-to-end distance analyses (Figures 6**l**,**m**): the control group maintained persistently low values, whereas ff19SB-PUD+ exhibited large fluctuations, consistent with the dynamic, flexible nature of IDPs.

Having established that ff19SB-PUD+ better captures IDP conformational behavior, we then examined whether this improvement extends to IDP-ligand binding. The multivalent and flexible binding interfaces of IDPs and intrinsically disordered regions (IDRs) enable them to serve as important ligand-binding domains, as previously demonstrated^27,28^. Here, we focused on the transactivation domain (TAD) of p53 and its interactions with two small-molecule inhibitors: PKU-MDL-RH 1047 and PKU-MDL-RH 1050 (hereafter referred to as “ligand 1047” and “ligand 1050”), both identified from the SPECS library using the IDP Drug Virtual Screening (IDPVDS) computational strategy^28^ and experimentally validated by surface plasmon resonance^28^. We conducted 3 μs simulations of the p53-TAD1-ligand complexes using ff19SB-PUD+ and ff19SB (as a control). The initial structures of these systems are shown in Figures 6**n** and 6**r**, respectively. A contact state was defined as any atom pair between p53-TAD1 and the ligand within a minimum distance of 6 Å, consistent with the previous work^27^, and the contact number per frame was taken as the total count of such atom pairs.

The minimum distance distributions (Figures 6**o**,**s**) revealed a higher peak below 6 Å for ff19SB-PUD+ compared to ff19SB, indicating a tighter p53-TAD1-ligand binding mode. Consistently, the total contact violin plots (Figures 6**p** and 6**t**) showed that ff19SB-PUD+ produced more frames with higher contact numbers, reflecting stronger and more extensive protein-ligand interactions. Residue-wise contact frequency analyses (Figures 6**q** and 6**u**) —defined as the fraction of contact-state frames over the total trajectory—revealed that both force fields displayed an elevated contact frequency in the central region around tryptophan W23, in agreement with previously reported binding patterns^27^. Notably, this shared feature validates the physical plausibility of both force fields, while the quantitative enhancements in ff19SB-PUD+ highlight its superior capacity to capture the full extent of IDP-ligand interactions. Taken together, these results demonstrate that ff19SB-PUD+ has substantially improved the simulation of IDP conformational dynamics and IDP-ligand binding, underscoring the value of PUD+’s extensive conformational sampling in force field reparameterization.

## Discussion

Accurate sampling of side-chain conformations is essential for faithfully capturing protein dynamics; however, existing datasets often suffer from insufficient backbone or side-chain coverage. To address this gap, we developed PUD+, that represents the largest DFT-level dataset for protein-related systems to date, offering a broader and more uniform coverage of the backbone and side-chain dihedral conformational space than existing biomolecule datasets and effectively bridging AI-driven simulation and classical simulations with higher accuracy and better fidelity. ViSNet-PIMA trained on PUD+ exhibited enhanced generalizability, with improvements evident not only in prediction accuracy for long-side-chain residues but also in the consistent performance gains aligned with increasing training set size. These findings indicate that data diversity and scale are key determinants of MLFF generalization. Furthermore, AI^2^BMD simulations demonstrated improved dynamic fidelity and simulation stability. Incorporating PUD+ into classical force field reparameterization (ff19SB-PUD+) also improved the simulation behavior of IDP systems, suggesting that high-accuracy, broad-coverage ab initio data holds substantial value for enhancing the physical realism of classical force fields.

Beyond the applications validated above, the potential value of PUD+ likely extends much further. Since the dataset simultaneously provides systematically sampled conformations, DFT energies, and atomic forces, its utility is not confined to a single level of force field development. A natural extension would be to use PUD+ for a more comprehensive parameterization of entire all-atom classical force fields (rather than solely CMAP term alone), and even for constructing coarse-grained force fields. This would enable a single, unified, high-accuracy, systematically sampled dataset to underpin multiscale dynamic simulations spanning from all-atom to coarse-grained representations, thereby mitigating the biases introduced by inconsistent parameter sources across different scales.

Nevertheless, we recognize that the conformational space generated through grid-based dihedral sampling combined with enhanced sampling strategies, however systematic, cannot achieve exhaustive coverage, particularly given the combinatorial explosion of side-chain degrees of freedom in high-dimensional space, where sparsely populated or boundary conformations may still remain elusive. Moreover, although force fields parameterized on PUD+ showed improvements for IDP systems, this may still be insufficient for protein classes with more distinctive structural or chemical environments, for example, transmembrane proteins embedded in lipid bilayers, or highly heterogeneous IDP families. For such scenarios, future work could pursue AI-driven sampling strategies tailored to specific systems, combined with active learning approaches that dynamically identify under-sampled conformational regions during simulation and supplement them accordingly. Such workflows could achieve a better balance between sampling efficiency and coverage completeness at a reduced computational cost.

While dipeptides facilitate the systematic enumeration of backbone and side-chain dihedral combinations, their local nature limits the model’s capacity to learn long-range interactions spanning multiple residues. Future studies should extend sampling to longer fragments, including tripeptides and tetrapeptides, to better capture long-range interactions. Additionally, given the important role of post-translational modifications (e.g., glycosylation, lipidation) in regulating protein function, incorporating carbohydrate and lipid molecules into the sampling scheme will extend the dataset’s coverage of the local chemical environments at modification sites, laying the groundwork for future MLFFs capable of handling post-translationally modified systems.

## Methods

### PUD+ dataset construction

The Protein Unit Dataset Plus (PUD+), comprising a 20 million backbone-sampling-enhanced (BB Sampling) portion and a 20 million side-chain-sampling-enhanced (SC sampling) portion. The original PUD dataset derived from AI^2^BMD serves as the BB sampling portion^6^. Briefly, PUD was constructed by performing a 2D grid scan of the backbone φ and ψ dihedral angles (−180° to 175°, in 5° interval) across 20 canonical dipeptides and the capped peptide ACE-NME, generating a set of anchor conformations. Each anchor underwent geometry optimization under the SMD solvent model, followed by quantum simulations of 225 fs per dipeptide anchor (2,025 fs for ACE-NME). Single point energies and atomic forces were subsequently recalculated without implicit solvent. All quantum mechanical calculations were carried out at the DFT level using the M06-2X functional with the 6-31G* basis set in ORCA 5.0.1, yielding approximately 20 million conformations with comprehensive backbone conformational coverage.

In this study, we generated the 20-million SC sampling portion with a newly designed pipeline: We prepared the initial topology and coordinate files for the 20 standard dipeptide types and the ACE-NME peptide using the ‘Sequence’ command in the tleap module of AmberTools20^29^. Each system then underwent energy minimization using the sander module of AMBER to relieve steric clashes and relax local geometries. The minimization followed a 1,000-cycle protocol under a generalized Born implicit solvent environment, transitioning from steepest descent to conjugate gradient method at the midpoint. Chirality restraints were applied throughout via the makeCHIR_RST script to preserve stereochemical integrity. The minimized structures then served as initial points for REMD^22^ simulations across four temperature replicas at 300 K, 380 K, 470 K, and 570 K. Each replica was first equilibrated for 200 ps under the Langevin thermostat, followed by a 10 ns production run. Electrostatic interactions were treated using the Generalized Born implicit solvent model (igb = 5)^30^ with no explicit periodic boundary conditions, and a long-range cutoff of 999 Å was applied to effectively eliminate cutoff-related artifacts for nonbonded interactions. Temperature exchanges between adjacent replicas were attempted every 2 ps, and a 2 fs timestep was maintained with bond constraints enforced by the SHAKE algorithm^31^. From the resulting REMD trajectories, we uniformly extracted 5,000 structures.

Then these structures were directly subjected them to 225 fs quantum simulations. The final 200 fs of each trajectory were retained as SC sampling dataset. All quantum simulations and single point energy calculations were performed at the DFT level using the M06-2X functional^23^ with the 6-31G* basis set in ORCA 5.0.1^32^. Tight convergence criteria were applied at each stage, and successful convergence was required before proceeding to the next step. Temperature was controlled by the Canonical Sampling through Velocity Rescaling (CSVR) thermostat at 290 K^33^, and the formal charge of each system was assigned based on summed residue charges at pH 7. The SC sampling calculations consumed approximately 12,465,271 CPU core hours (∼1,422 CPU core years) in total, yielding 20 million DFT-level conformations with substantially enhanced side-chain diversity.

We then calculated the backbone dihedral angles φ and ψ, as well as side-chain dihedral angles (excluding Glycine), for both the BB sampling and SC sampling parts. To provide a general overview of conformational sampling, Ramachandran plots were generated for all dipeptides (Figure 2**a-c**), and 2D distribution plots were constructed for all dipeptides possessing both χ1 and χ2 dihedral angles (Figure 2**d-f**). For residues with only χ1, the distribution of this dihedral was plotted separately (see Supplementary Information Figure 2). Individual Ramachandran plots and 2D distribution plots for each dipeptide are provided in the Supplementary Information. For comparison, equivalent plots were also generated for the Dunbrack raw dataset (Figure 2**g** and 2**i**) and MEDFORD (Figure 2**h** and 2**j**). As the standard Dunbrack rotamer library is not well-suited for 2D dihedral distribution analysis due to its limited data points and discrete binning strategy, the Dunbrack raw dataset was used in its place. All 2D dihedral distribution plots were generated using a 3° bin width, except for the MEDFORD Ramachandran plot, which was binned at 10° to match its original data resolution. For structural clustering, the φ, ψ, and χ1 conformational space of PUD+ was partitioned into nine clusters using k-means clustering. Conformations from the BB sampling portion, SC sampling portion, and the Dunbrack raw dataset were then mapped onto these nine clusters, and the structure closest to each cluster center was selected as the representative conformation.

### MLFF settings and training details

The 40 million PUD+ dataset was partitioned into training, validation, and test sets following a standardized pipeline. For each dipeptide (including ACE-NME), 5% of the data were uniformly sampled from both the BB sampling and SC sampling portions to form the independent test set. The remaining data for each dipeptide were then split into training and validation sets at an 8:2 ratio. The final PUD+ training and validation sets were assembled by combining the respective subsets from both the BB sampling and SC sampling portions across all dipeptides. The PUD dataset, containing only the BB sampling part, was retained as a control.

For model training, ViSNet-PIMA^25^ was trained on 0.5%, 5%, 20%, and 100% of the PUD+ training set. As a control, ViSNet-PIMA was also trained on 40% of the PUD training set, which contained an equal number of samples as the 20% PUD+ training set. The model trained on the full (100%) PUD+ used eight ViSNet layers and one PIMA layer, while all other models (0.5%, 5%, and 20% PUD+, as well as the 40% PUD control) used four ViSNet layers and one PIMA layer; all models shared 128-dimensional node and edge embeddings, and angular information was captured via spherical harmonic expansions up to degree l_max_ = 1. To build the radius graph, a neighbor cutoff radius of 15 Å was applied across all systems. The training objective combined energy and force prediction into a single mean-squared-error loss, with energy and force terms weighted at a ratio of 1:50 (approximately 0.02 and 0.98, respectively). Parameters were optimized using the AdamW optimizer built on an AMSGrad base transform, with gradients clipped to a maximum norm of 500 and no additional weight decay. The learning rate followed a warm-up schedule, increasing linearly from 1 × 10^-4^ to a peak of 5 × 10^-4^, followed by a linear decay to a final value of 1 × 10^-6^. Training used a batch size of 64 or 128, depending on the size of the respective training subsets. Prior to training, element-specific atomic reference energies were subtracted from each system’s total energy. In addition, we trained several state-of-the-art MLFFs, including MACE^34^, AIMNet2^35^, SO3krates^36^, Equiformer V2^37^, and ViSNet^38^, on 5% of the PUD+ training set. Default hyperparameter settings were applied for most models; for models with exceptionally large parameter sizes, the model capacity was reduced to a level comparable to the other models to ensure a fair comparison. All models were trained on an NVIDIA GeForce RTX 5090 GPU. Detailed training hyperparameters are provided in the Supplementary Table S6.

### AI^2^BMD simulations

For the Ace-Ala_15_-Nme simulation, we first generated a fully extended structure using PyMOL^39^. We then conducted a 250 ps AI^2^BMD simulation in vacuum at 300 K under NVT ensemble, employing periodic boundary conditions (PBC) with the particle mesh Ewald (PME) method for electrostatic interactions. The fragmentation strategy was employed, and the ViSNet-PIMA model trained on the full PUD+ training set was utilized to predict the potential energies and atomic forces of the fragmented dipeptides. Then we sampled frames every 10 ps and calculated the potential energies and atomic forces at DFT level using ORCA 6.1.1, with the M06-2X functional and the 6-31G* basis set consistent with the training data. The energies and atomic forces of the snapshots sampled during the AI^2^BMD simulation were calculated by ViSNet-PIMA trained on PUD+. For comparison, we also calculated these values by Amber ff19SB force field^3^. The potential energy of the first frame was subtracted to yield the relative potential energies for comparison.

For the STE7 docking motif (sequence RRNLKGLNLNLH, PDB ID: 2B9H), we first generated a fully extended structure using PyMOL and then conducted a 1 ns AI^2^BMD simulation in vacuum at 300 K under NVT ensemble to obtain a compactly folded initial conformation. Topology and coordinate files for the folded structure were then prepared using the tleap module of AmberTools20 under the ff19SB force field. The system then underwent energy minimization using the sander module of AMBER under a Generalized Born implicit solvent environment (igb=5), following a 1,000-cycle protocol that transitioned from steepest descent to conjugate gradient at the midpoint. Chirality restraints were applied throughout via the makeCHIR_RST script to preserve stereochemical integrity. The minimized structures served as starting points for REMD simulations conducted under the same force field and solvent settings. Eight temperature replicas were deployed at 300, 400, 500, 600, 700, 800, 900, and 1000 K. Each replica was first equilibrated for 200 ps under the Langevin thermostat (collision frequency 1.0 ps^-^¹), followed by a 10 ns production run with exchange attempts between adjacent replicas every 2 ps. A 2 fs timestep was maintained throughout, with bond constraints enforced by the SHAKE algorithm. Trajectories from all eight replicas were combined and aligned using CPPTRAJ. K-means clustering was then applied to the full trajectory based on backbone RMSD against the initial structure, yielding 10 clusters in total. The structure closest to each cluster center was selected as the representative conformation. These 10 representative structures were subsequently classified into folded (Rg < 8.0 Å), intermediate (8.0 Å ≤ Rg ≤ 10.0 Å), and unfolded (Rg > 10.0 Å) states based on their radius of gyration (Rg), resulting in three folded, four intermediate, and three unfolded structures for subsequent simulations.

Prior to the AI^2^BMD production simulations, each initial structure was solvated in a cubic TIP3P water box with a minimum solvent buffer of 20 Å from the protein surface, using the Amber ff19SB force field for the protein and the TIP3P force field for water. Periodic boundary conditions (PBC) were applied throughout, with long-range electrostatic interactions treated using the particle mesh Ewald (PME) method. The system was neutralized and supplemented with Na^+^/Cl^-^ counterions, parameterized according to the Joung-Cheatham ion model optimized for TIP3P water, to approximate physiological ionic strength. Energy minimization was first performed using a 100-cycle protocol (maxcyc=100, ncyc=50), transitioning from steepest descent to conjugate gradient at the midpoint, to remove steric clashes and relax the initial geometries. The minimized structures then underwent a stepwise equilibration protocol. First, the system was gradually heated from 0 K to 300 K over 40 ps under NVT ensemble using the Langevin thermostat, with positional restraints applied to all protein residues to prevent structural distortion during heating. This was followed by three successive NVT equilibration stages (20 ps each) with progressively relaxed restraints: all protein residues were restrained in the first stage to allow solvent relaxation around the fixed protein, only Cα atoms were restrained in the second stage to permit side-chain flexibility, and all restraints were removed in the third stage to allow full system relaxation. Finally, two NPT equilibration stages were performed to stabilize the system pressure and density: a short 10 ps stage to allow initial volume adjustment, followed by a longer 100 ps stage to ensure full convergence of the system density. From each preprocessed structure, we then conducted 1 ns AI^2^BMD simulations using ViSNet-PIMA trained on PUD and PUD+, respectively, at 300 K with AMOEBA polarizable water model under NVT ensemble. Trajectories were sampled every 100 ps; for collapsed trajectories, only frames prior to the collapse point were retained. To directly evaluate the accuracy of ViSNet-PIMA trained on dipeptide data, each polypeptide was fragmented into dipeptide and ACE-NME units. DFT-level potential energies and atomic forces were then computed for these fragments using ORCA 6.1.1 with the M06-2X functional and the 6-31G* basis set, consistent with the training data. Absolute errors in potential energies and atomic forces were calculated against the DFT reference values. The same set of sampled frames was used for residue-wise, per-atom MAE analysis of both potential energies and atomic forces.

### CMAP reparameterization in Amber ff19SB force field

For the CMAP reparameterization, dipeptide samples in PUD+ were filtered by their rotamer states, retaining only those belonging to the most prevalent rotamer for each residue type. This selection prevented side-chain rotamer energy differences from being introduced into the backbone correction map. For each grid point of the 24 × 24 CMAP, the 10 nearest data points were aggregated, and their average value was assigned to the corresponding grid point. DFT-level potential energies were taken directly from the dataset labels, calculated using the M06-2X functional with the 6-31G* basis set, while MM potential energies were computed using ff19SB without the CMAP term. Both DFT and MM potential energies were referenced to their respective minima to obtain relative potential energies. The CMAP correction term U_cmap_ was then derived according to the following formula:

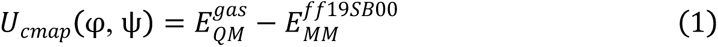

where 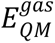 denotes the gas-phase DFT-level potential energy and 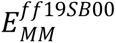 denotes the molecular mechanics potential energy calculated by the ff19SB force field with the CMAP correction term set to zero (hence “00”). The newly generated CMAP was used to replace the original CMAP in ff19SB. Implementation within the AMBER software suite is natively supported through the bicubic spline interpolation function.

### Classical MD simulation for IDPs

We conducted classical MD simulations with PUD+ reparameterized CMAP term on two intrinsically disordered protein (IDP) systems. For the Fused in sarcoma (FUS) protein, an IDP with 526 residues, we obtained the initial structure from Kumar Sarthak, *el al*^21^. The initial structure was derived from an equilibrated coarse graining simulation^40^. We prepared the system under both ff19SB and ff19SB-PUD+ (where the original CMAP was replaced with PUD+ reparameterized CMAP) force fields, with the FUS protein solvated in an octahedral box of SPC/E water with a minimum padding of 11 Å. Potassium (K⁺) and chloride (Cl⁻) ions were added to neutralize the system and approximate physiological ionic conditions. Each system underwent a two-stage energy minimization. In the first stage, 1,000 cycles of minimization (ncyc=500, transitioning from steepest descent to conjugate gradient at the midpoint) were performed with harmonic positional restraints (restraint weight of 500 kcal/mol·Å²) applied to all non-hydrogen protein atoms, allowing the solvent to relax while preserving the initial protein conformation. In the second stage, all restraints were removed and an additional 1,000 cycles of unrestrained minimization were carried out to relax the entire system. The system was then gradually heated from 0 K to 300 K over 20 ps under NVT conditions using the weak-coupling thermostat, with restraints (restraint weight of 10 kcal/mol·Å²) maintained on all non-hydrogen protein atoms to prevent structural distortion during heating. Subsequent simulations were performed under periodic boundary conditions (PBC), with long-range electrostatic interactions treated using the particle mesh Ewald (PME) method and a real-space cutoff of 9 Å for short-range non-bonded interactions. The system was next equilibrated for 1 ns under NPT conditions at 300 K and 1 bar using the Monte Carlo barostat to stabilize the system density. Bond lengths involving hydrogen atoms were constrained using the SHAKE algorithm throughout, with a 2 fs timestep. Production simulations of 1,500 ns were then conducted under the same NPT conditions for both the ff19SB and ff19SB-PUD+ systems. The CPPTRAJ module of Amber was used to calculate the radius of gyration (Rg) and the end-to-end distance.

For the IDP-ligand binding system, we selected the important tumor suppressor p53, which contains a 39-residue IDP region at N-terminus, the transactivation domain (TAD1, PDB ID: 2K8F). Previous studies have shown that TAD1 binds to two small molecules, the ligand PKU-MDL-RH 1047 (C_22_H_16_N_3_OS_2_^+^, denoted as 1047) and PKU-MDL-RH 1050 (C_23_H_16_NOS_2_^+^, denoted as 1050) from the SPECS library^28^. For p53-TAD1-1047 and p53-TAD1-1050 systems, the protein was described by the ff19SB-PUD+ (with ff19SB serving as a control) force field, while the two small-molecule ligands (1047 and 1050) were parameterized using the GAFF2 force field with partial charges derived from the respective mol2 files. Each complex was solvated in a rectangular box of TIP4P-D water with a minimum padding of 15 Å, and K⁺ and Cl⁻ ions were added to neutralize the system. Each system underwent a two-stage energy minimization following the same protocol described above, with harmonic positional restraints (500 kcal/mol·Å² for the first stage, then removed for the second) applied to all non-hydrogen protein and ligand atoms. The minimized system was then heated from 0 K to 298 K over 20 ps under NVT conditions using the weak-coupling thermostat, with positional restraints (restraint weight of 10 kcal/mol·Å²) maintained on all non-hydrogen protein and ligand atoms. As above, periodic boundary conditions were applied throughout the subsequent simulations, with long-range electrostatics treated via the particle mesh Ewald method and a real-space cutoff of 8 Å. Equilibration was conducted in two NPT stages at 298 K and 1 atm using the Monte Carlo barostat: a short 10 ps stage for initial pressure stabilization, followed by a 1 ns stage to achieve full convergence of system density. Bond constraints and integration timestep followed the same SHAKE/2 fs setup described previously. Production simulations of 3 μs were then performed under the same NPT conditions for both the ff19SB and ff19SB-PUD+ systems. Ramachandran plots were generated from the ff19SB and ff19SB-PUD+ trajectories, computing backbone φ and ψ dihedral angles for all protein residues across every frame and binning them into two-dimensional histograms (180 × 180 bins, -180° to 180°) visualized on a logarithmic color scale. Four canonical secondary structure regions were annotated on the φ/ψ maps: α-helix (right-handed), φ ∈ [-160°, -20°], ψ ∈ [-120°, 60°]; β-sheet, φ ∈ [-180°, -40°], ψ ∈ [80°, 180°]; polyproline II (PPII), φ ∈ [-100°, -40°], ψ ∈ [100°, 180°]; and α-helix (left-handed), φ ∈ [20°, 100°], ψ ∈ [20°, 120°]. A zoomed-in region spanning φ ∈ [-40°, 30°] and ψ ∈ [60°, 130°], corresponding to the transition zone between the PPII and left-handed α-helical basins, was additionally visualized using finer binning (60 × 60 bins) to resolve conformational shifts within this range.

## Author contributions

T. W. led, conceived, designed and supervised this study. D. W. conducted experiments and analysis. D. W. and T. W. wrote the original manuscript. All authors approved the final version of manuscript.

