## Supplementary Information for "Ab initio side-chain sampling with PUD+ enables high-fidelity protein dynamics across AI-driven and classical simulations"

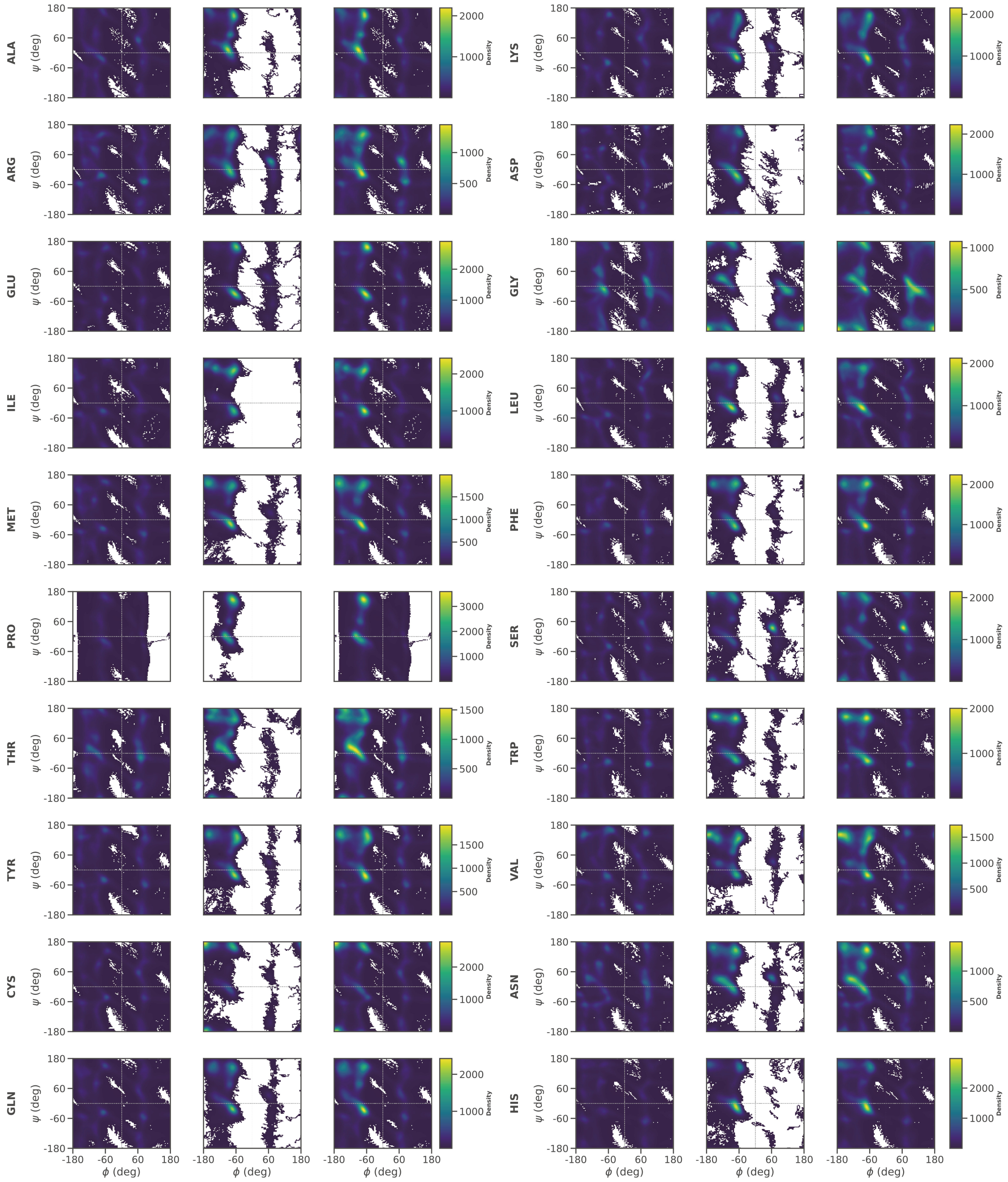

**Supplementary Figure 1. Main-chain conformational space coverage for each dipeptide type.** For each dipeptide type, Ramachandran plots of the main-chain dihedral angles (φ, ψ) are shown for BB sampling (left), SC sampling (middle), and the complete PUD+ dataset (right).

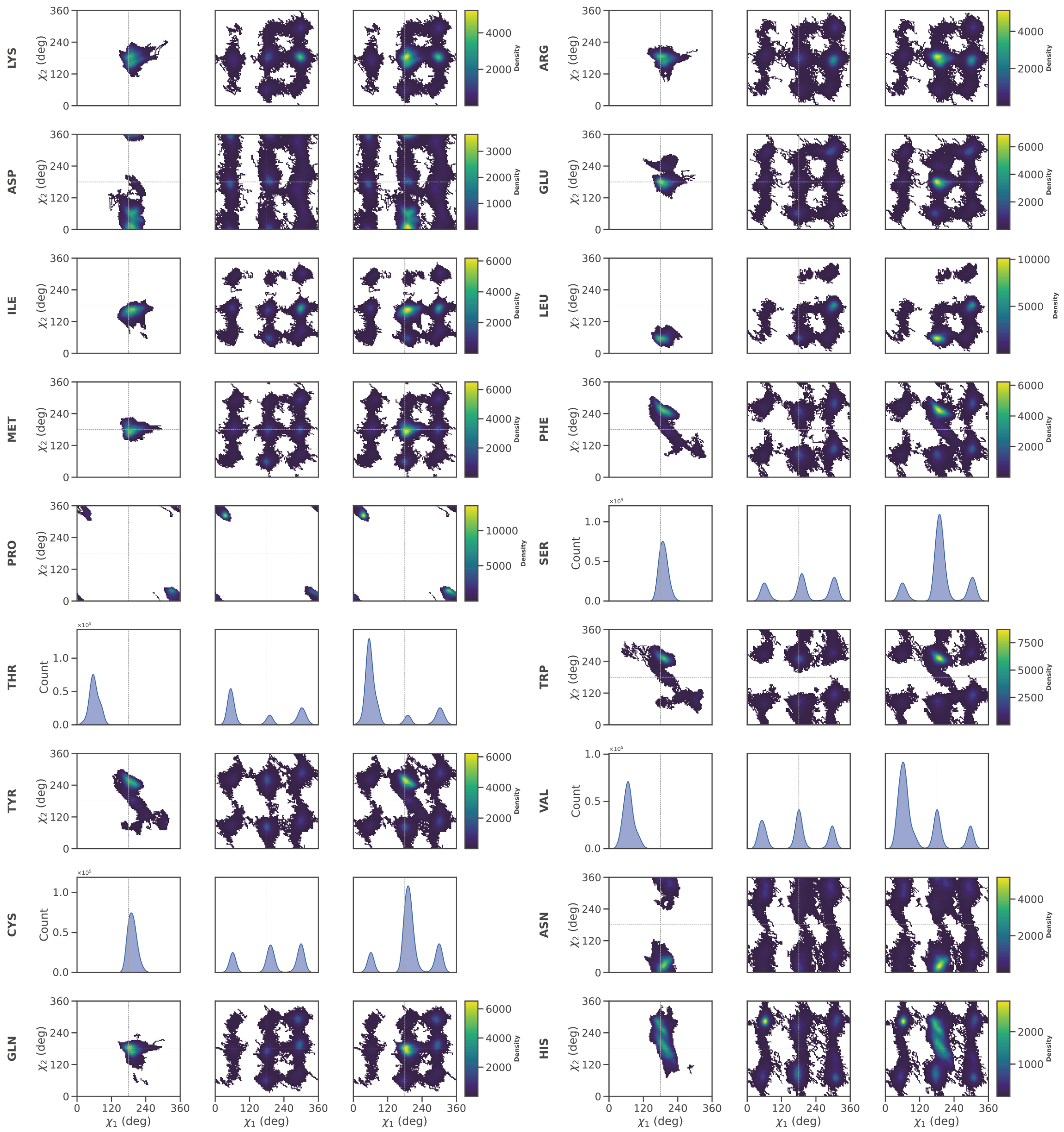

**Supplementary Figure 2. Side-chain conformational space coverage for each dipeptide type.** For each dipeptide type, the distributions of side-chain dihedral angles (χ1, χ2) are shown for BB sampling (left), SC sampling (middle), and the complete PUD+ dataset (right). Dipeptides possessing only a single side-chain dihedral angle (χ1) are displayed in 1D χ1 distribution plots. Alanine and glycine are excluded due to the absence of a χ1 dihedral angle.

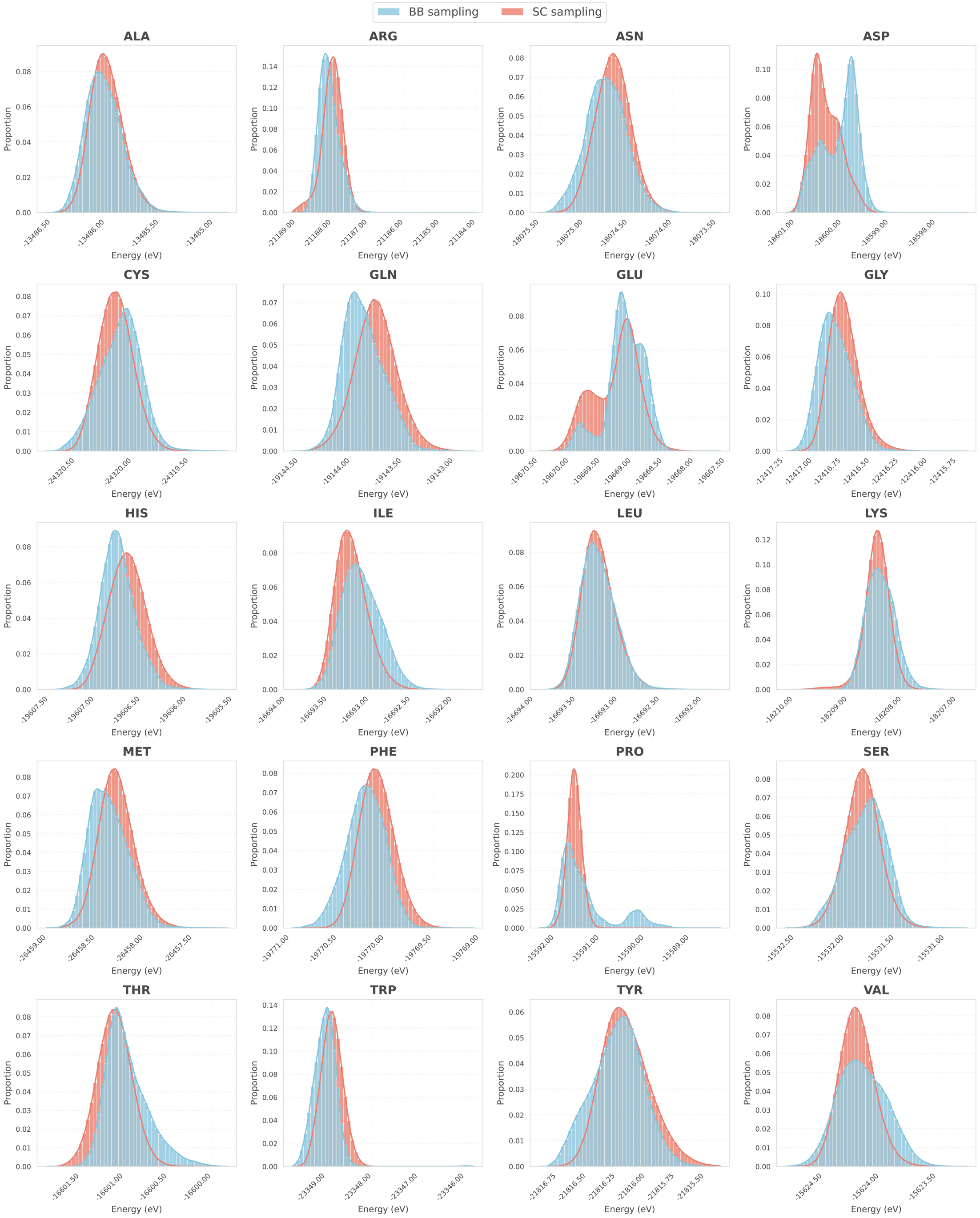

**Supplementary Figure 3. Distributions of potential energies for each dipeptide type.** Distributions for BB sampling and SC sampling are shown in skyblue and salmon, respectively.

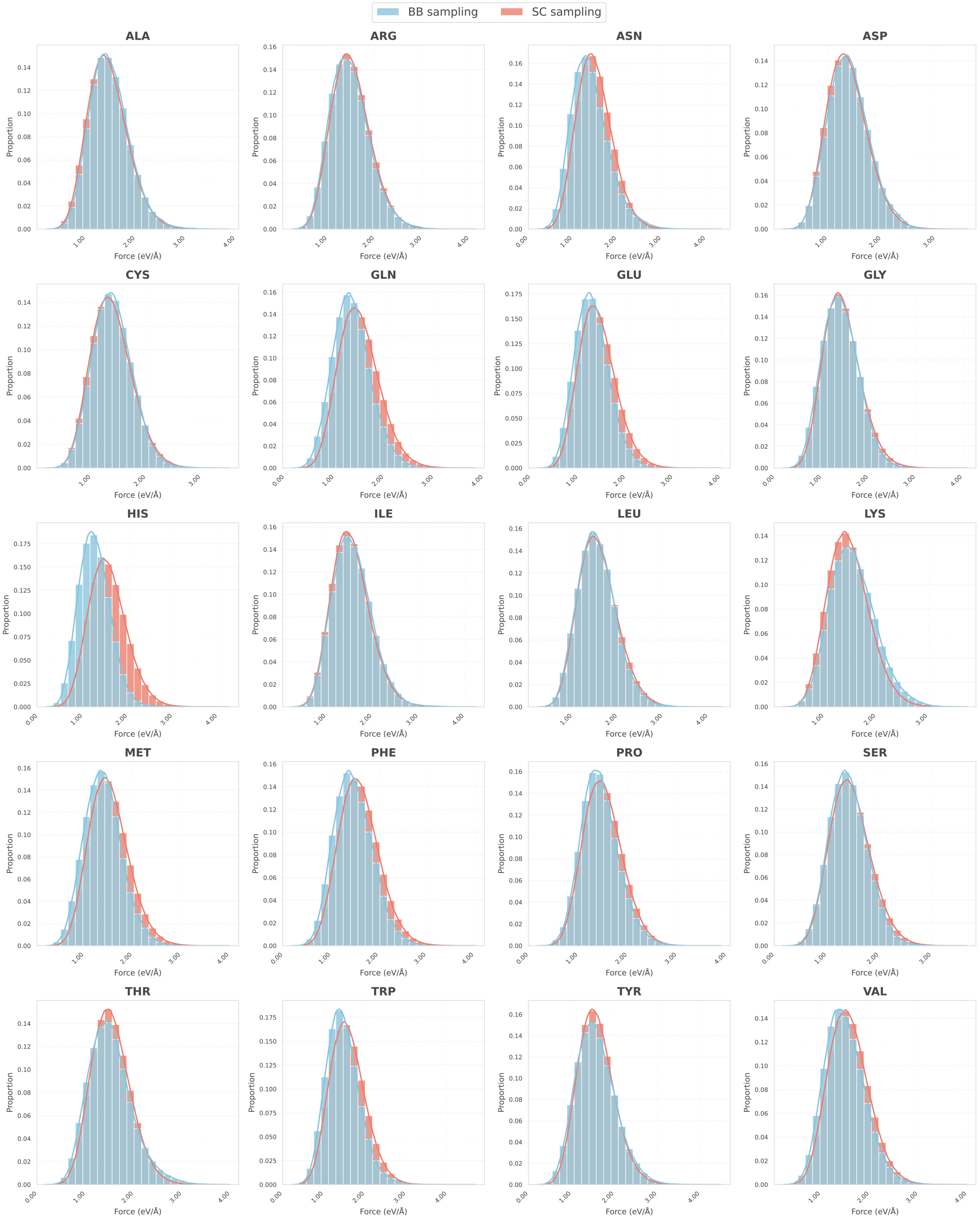

**Supplementary Figure 4. Distributions of atomic forces on main-chain atoms for each dipeptide type.** Distributions for BB sampling and SC sampling are shown in skyblue and salmon, respectively. The main-chain atomic force is defined as the average force norm of all backbone heavy atoms within each dipeptide conformation.

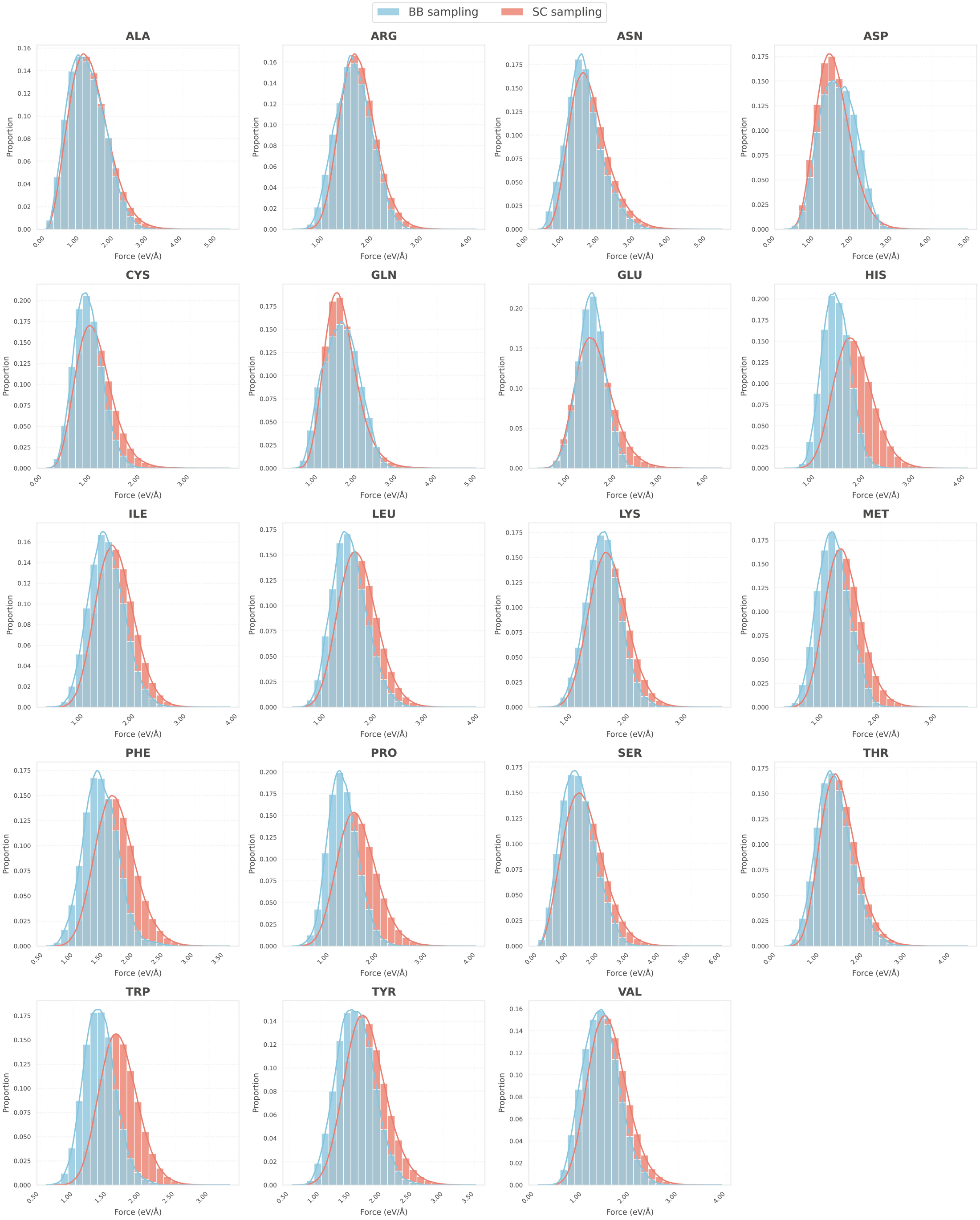

**Supplementary Figure 5. Distributions of atomic forces on side-chain atoms for each dipeptide type.** Distributions for BB sampling and SC sampling are shown in skyblue and salmon, respectively. The side-chain atomic force is defined as the average force norm of all heavy atoms on the side-chain within each dipeptide conformation.

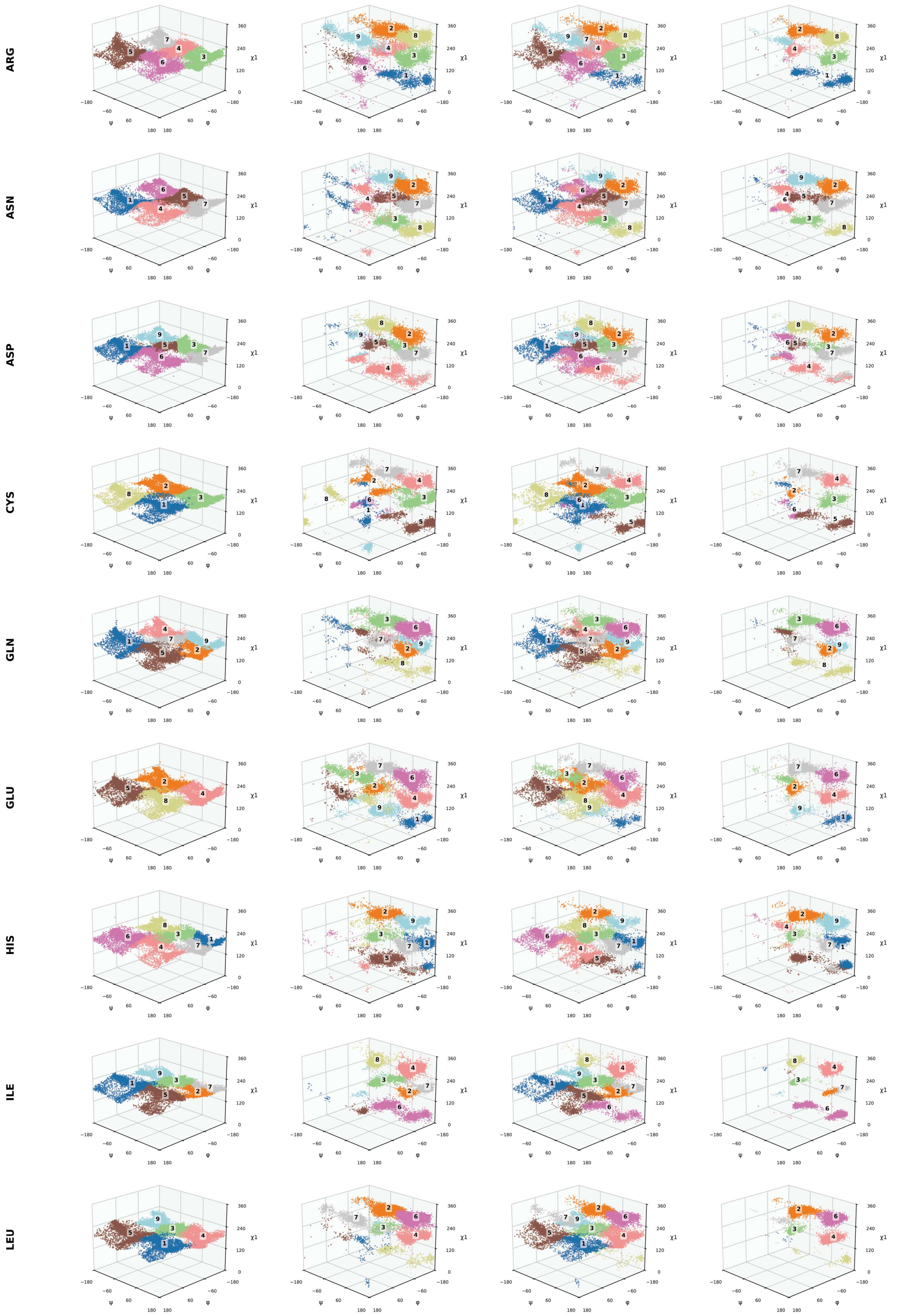

**Supplementary Figure 6. Structural clustering analysis for each dipeptide type.** For each dipeptide type, structural clustering results are shown for BB sampling (first column), SC sampling (second column), PUD+ (third column) and Dunbrack raw dataset (fourth column). The conformational space is defined in three dimensions

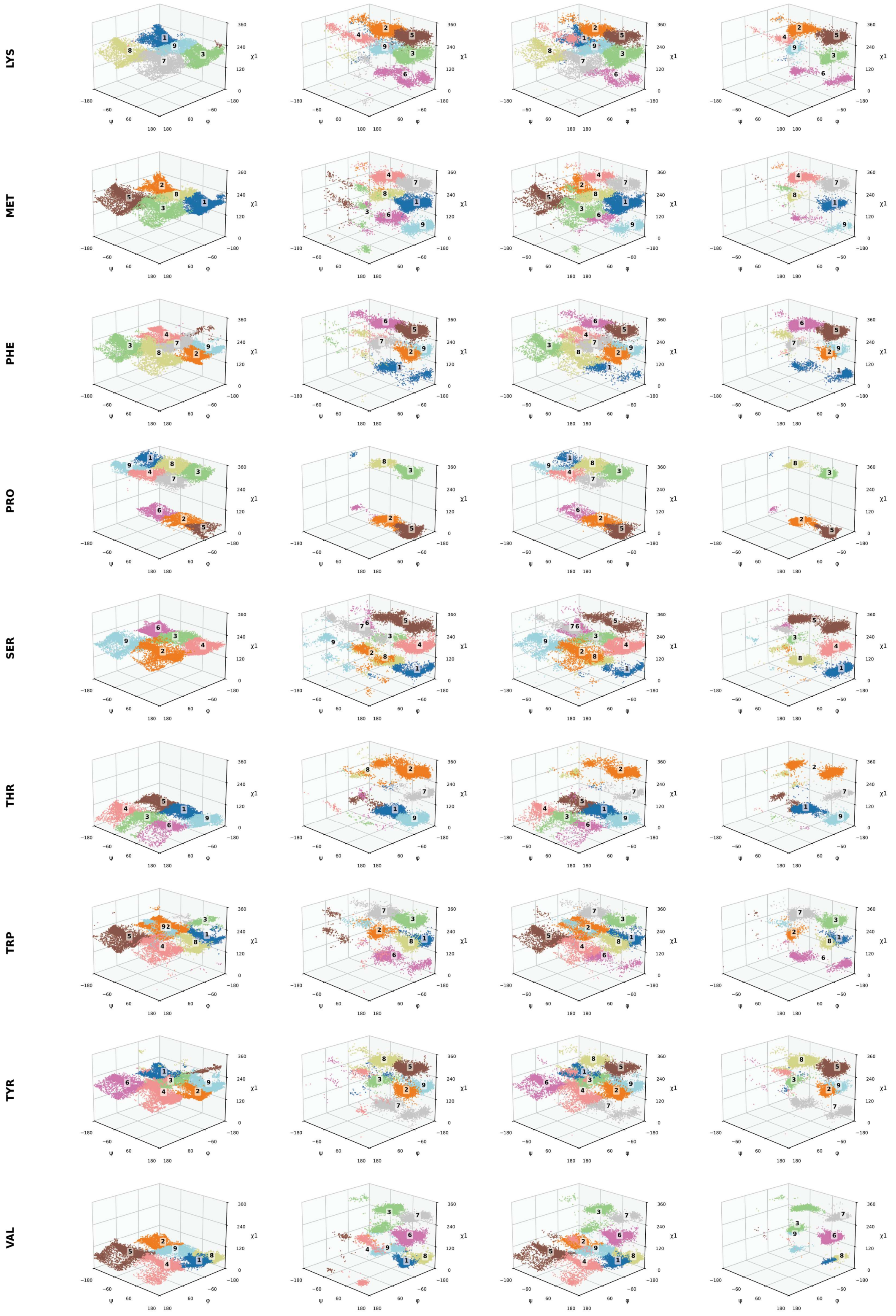

using the backbone dihedral angles (φ and ψ), and the first side-chain dihedral angle (χ1). Alanine and glycine are excluded due to the absence of a χ1 dihedral angle. For each dipeptide type, nine clusters were identified based on all conformations in PUD+ and visualized by distinct colors.

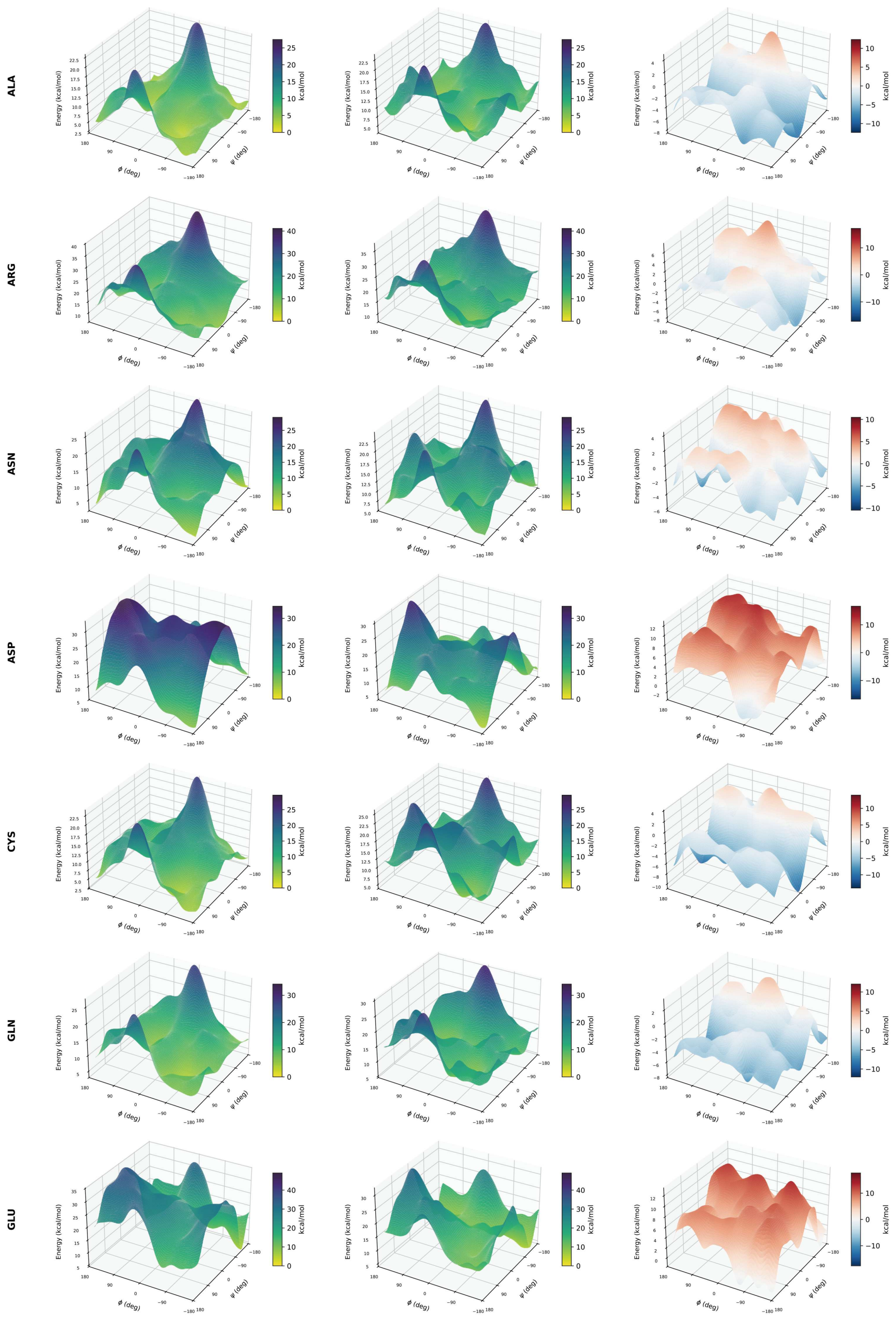
ALA
ARG
ASN
ASP
CYS
GLN
GLU
Energy (kcal/mol)
ϕ (deg)
ψ (deg)
kcal/mol

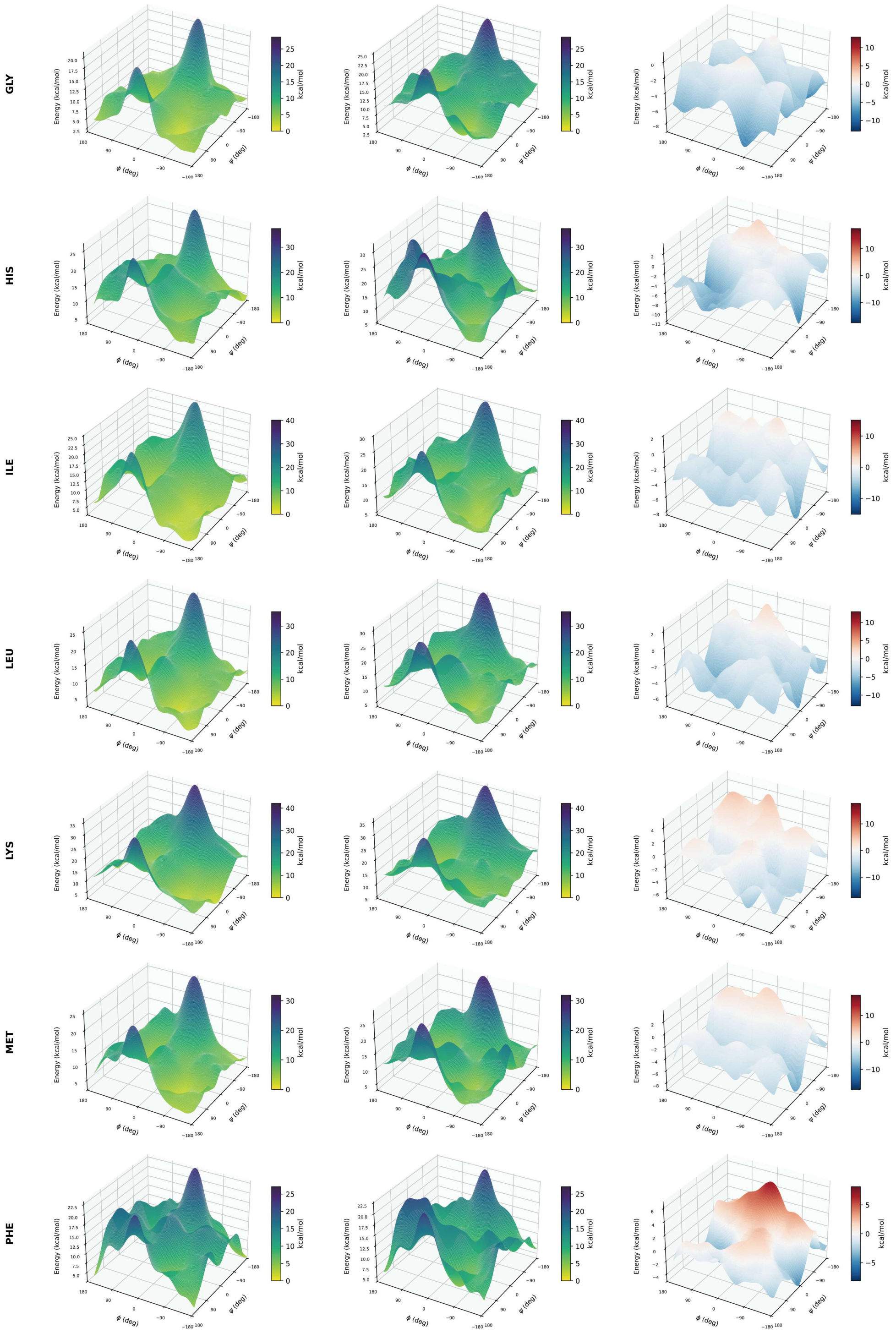
GLY
HIS
ILE
LEU
LYS
MET
PHE
Energy (kcal/mol)
ϕ (deg)
ψ (deg)
kcal/mol

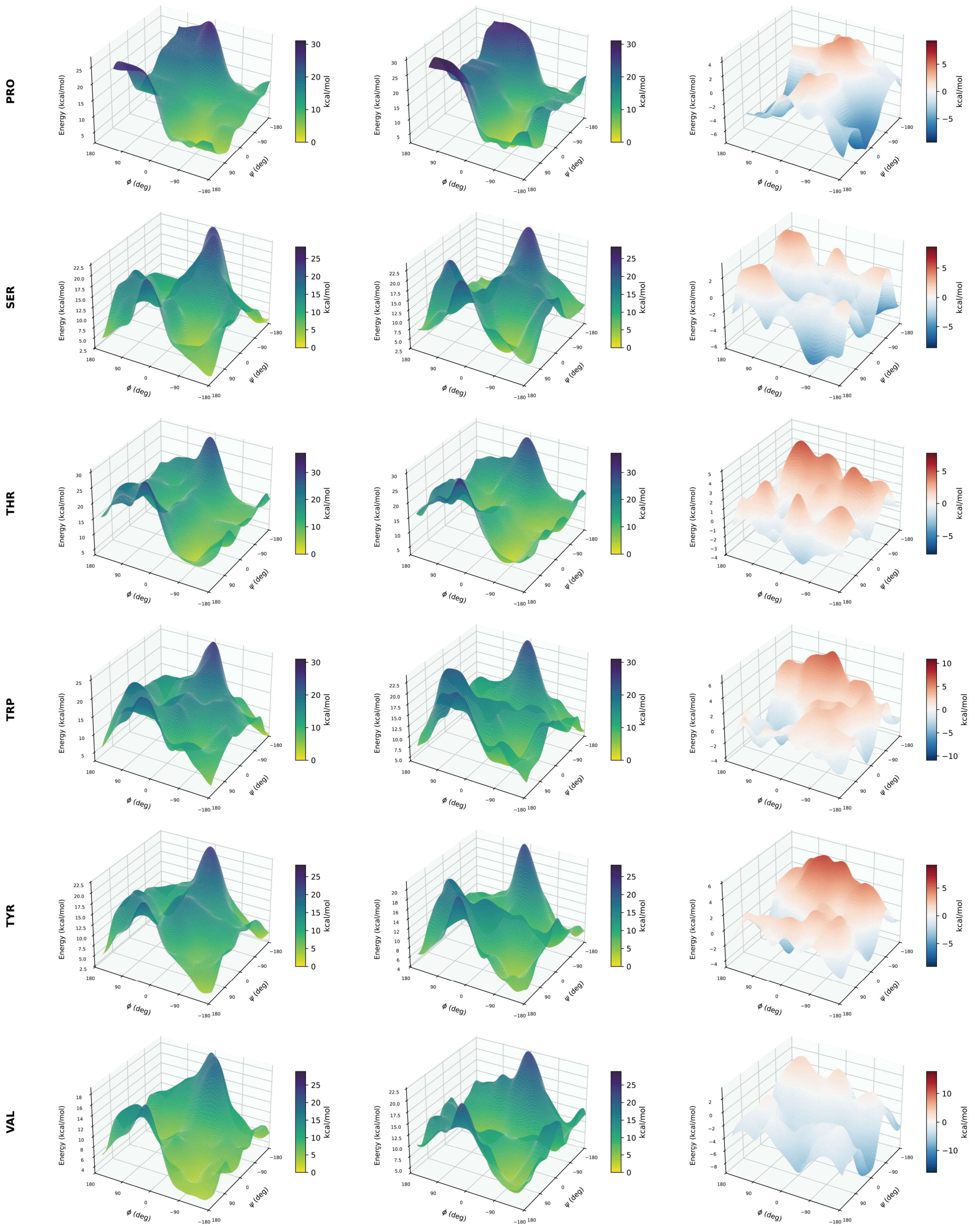

**Supplementary Figure 7. Three-dimensional (3D) plots of the relative potential energy surfaces (PESs) for each dipeptide type.** From left to right, the panels show the DFT relative PES, the MM relative PES and the difference between the two surfaces. Both DFT and MM potential energies were referenced to their respective minima to obtain relative potential energies.

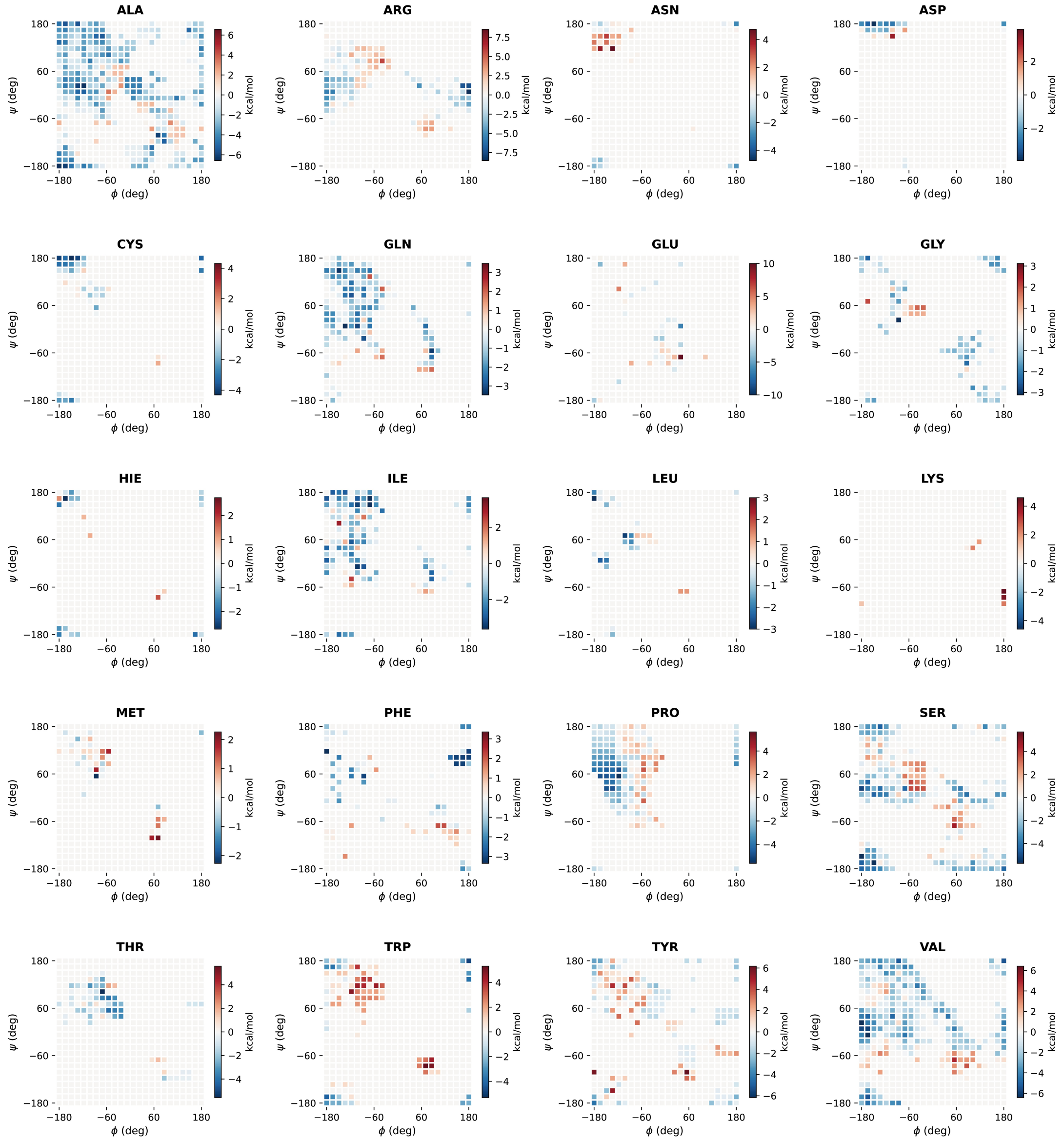

**Supplementary Figure 8. Two-dimensional (2D) plots for CMAP potential energy term constructed using PUD+.** The CMAP potential energy term $U_{CMAP}$ is defined as the difference between the DFT and MM relative potential energy surfaces. For each dipeptide type, a 24 × 24 grid was applied, and the 10 nearest data points were aggregated, with their average value assigned to the corresponding grid point.

**Supplementary Table 1. MAEs of energy and force predictions for ViSNet-PIMA models trained on PUD or PUD+, respectively.** Mean absolute errors of potential energies ($kcal \cdot mol^{-1}$ per atom) and atomic forces ($kcal \cdot mol^{-1} \cdot Å^{-1}$) on the PUD+ independent test set. The predictions were generated by ViSNet-PIMA trained on 40% PUD and 20% PUD+, respectively.

| Training Set | Energy | Force |
|---|---|---|
| PUD | 0.01414 | 0.1825 |
| PUD+ | 0.002576 | 0.05984 |

**Supplementary Table 2. MAEs of energy predictions across 20 kinds of dipeptides.** Mean absolute errors of potential energies (kcal·mol$^{-1}$ per atom) across 20 kinds of dipeptides on the PUD+ independent test set. The predictions were generated by ViSNet-PIMA trained on 40% PUD and 20% PUD+, respectively.

| Dipeptide Type | PUD | PUD+ |
|---|---|---|
| ALA | 0.001418 | 0.001755 |
| CYS | 0.003454 | 0.002347 |
| ASP | 0.003035 | 0.002585 |
| GLU | 0.004246 | 0.002211 |
| PHE | 0.002001 | 0.001796 |
| GLY | 0.001512 | 0.002312 |
| HIS | 0.002948 | 0.002084 |
| ILE | 0.003055 | 0.002561 |
| LYS | 0.003085 | 0.002130 |
| LEU | 0.002373 | 0.002031 |
| MET | 0.003673 | 0.001996 |
| ASN | 0.003200 | 0.001997 |
| PRO | 0.001662 | 0.001495 |
| GLN | 0.002532 | 0.002898 |
| ARG | 0.003876 | 0.002267 |
| SER | 0.002505 | 0.002003 |
| THR | 0.002577 | 0.001895 |
| VAL | 0.002478 | 0.001480 |
| TRP | 0.002278 | 0.002515 |
| TYR | 0.001955 | 0.001881 |

**Supplementary Table 3. MAEs of force predictions across 20 kinds of dipeptides.** Mean absolute errors of atomic forces (kcal·mol$^{-1}$·Å$^{-1}$) across 20 kinds of dipeptide on the PUD+ independent test set. The predictions were generated by ViSNet-PIMA trained on 40% PUD and 20% PUD+, respectively.

| **Dipeptide Type** | **PUD** | **PUD+** |
|---|---|---|
| ALA | 0.049730 | 0.040782 |
| CYS | 0.064310 | 0.052184 |
| ASP | 0.076995 | 0.061833 |
| GLU | 0.075724 | 0.064946 |
| PHE | 0.064339 | 0.052447 |
| GLY | 0.047613 | 0.040288 |
| HIS | 0.076604 | 0.061040 |
| ILE | 0.075490 | 0.060654 |
| LYS | 0.063749 | 0.053637 |
| LEU | 0.065812 | 0.053683 |
| MET | 0.061309 | 0.052941 |
| ASN | 0.071461 | 0.058232 |
| PRO | 0.098409 | 0.061000 |
| GLN | 0.060353 | 0.051702 |
| ARG | 0.109895 | 0.063850 |
| SER | 0.065503 | 0.053303 |
| THR | 0.087092 | 0.060730 |
| VAL | 0.067269 | 0.053015 |
| TRP | 0.081945 | 0.063395 |
| TYR | 0.068642 | 0.054401 |

**Supplementary Table 4. MAEs of energy and force predictions for ViSNet-PIMA models trained on PUD+ of varying scales.** Mean absolute errors of potential energies (kcal · $mol^{-1}$ per atom) and atomic forces (kcal · $mol^{-1}$ · $Å^{-1}$) on the PUD+ independent test set. The predictions were generated by ViSNet-PIMA trained on training subsets of varying scales.

| Subset Scale | Energy | Force |
|---|---|---|
| 0.5% | 0.01075 | 0.2081 |
| 5% | 0.004107 | 0.08977 |
| 20% | 0.002576 | 0.05984 |
| 100% | 0.001850 | 0.05024 |

**Supplementary Table 5. MAEs of energy and force predictions for different models trained on PUD+.** Mean absolute errors of potential energies (kcal·mol$^{-1}$ per atom) and atomic forces (kcal·mol$^{-1}$·Å$^{-1}$) on the PUD+ independent test set. The predictions were generated by various machine learning force fields trained on 5% of the PUD+ training set. The best results in each category are highlighted in bold.

| Model | Energy | Force |
|---|---|---|
| Mace | 0.01881 | 0.3166 |
| AIMNet2 | 0.008877 | 0.3084 |
| SO3krates | 0.005923 | 0.1917 |
| ViSNet | 0.004524 | 0.09060 |
| ViSNet-PIMA | **0.004107** | 0.08977 |
| Equiformer V2 | 0.006723 | **0.06550** |

**Supplementary Table 6. Hyperparameters for the model training of both ViSNet and ViSNet-PIMA.**

| Hyperparameters | Search space |
|---|---|
| Batch size | 64, 128 |
| Hidden size | 128 |
| Cutoff distance | 15.0 |
| Initial learning rate | 1e-4 |
| Peak learning rate | 5e-4 |
| Final learning rate | 1e-6 |
